# Codon usage determines tRNA-modification requirements for bacterial virulence

**DOI:** 10.64898/2026.09.09.750475

**Authors:** Frédéric Goormaghtigh, Minia Antelo Varela, Dirk Bumann

## Abstract

Bacterial virulence genes exhibit distinctive codon usage that may impose tRNA-modification requirements for efficient translation, but a comprehensive understanding is lacking. Here, a curated codon-tRNA modification map for *Salmonella* predicted six modifications that could preferentially support virulence gene translation. Mutant phenotypes in infected mice confirmed requirements for i⁶A_37_ and C^5^ modification of U_34_, whereas essential k^2^C_34_ could not be tested. Q_34_, Cm_32_/Um_32_, and Cm_34_ had limited impact consistent with weak decoding effects in other systems. Non-predicted tRNA modifications were not required. Synonymous reporter variants, proteomics, and targeted recoding demonstrated that virulence-associated codons directly conferred modification dependence. Additionally, four Leu-TTA codons in *ssrB*, the master regulator of *Salmonella* pathogenicity island 2 (SPI-2), contributed to MiaA-dependent SPI-2 gene expression. Extending our analysis to other pathogens predicted shared and divergent virulence dependencies consistent with published mutant phenotypes. Thus, codon usage explains tRNA-modification requirements for bacterial virulence.

## Main

Synonymous codons encode the same amino acid but are decoded by distinct or partially overlapping tRNA species that differ in abundance, charging and decoding properties. Modifications in the anticodon stem-loop (ASL) further shape these properties. At wobble position 34, modifications can expand or restrict codon recognition^1–5^, whereas modifications adjacent to the anticodon, at position 37, can stabilize codon–anticodon interactions and support accurate decoding^6,7^. Loss of these modifications can disrupt translation and proteostasis, but individual pathways can also have more selective effects on stress adaptation and infection^8–18^.

Individual pathways have well-documented, gene-specific consequences. In *E. coli*, MiaA-dependent i^6^A_37_ is required for efficient translation of the general stress regulator *rpoS*, whose transcript is enriched in Leu-TTA/TTG codons^9^. MnmEG-dependent modification of U_34_ supports expression of *Shigella* virulence-plasmid genes that are enriched in modification-dependent codons^10^, and these and other tRNA-modifying enzymes are required for virulence in diverse pathogens^19,20^. Why particular tRNA modifications matter during infection, and why others do not, nonetheless remains unclear, because a comprehensive analysis of codons requiring tRNA modifications in virulence-associated genes is lacking. Moreover, the interpretation of individual mutants can be complicated by polar effects of the mutation on downstream genes and unrelated moonlighting effects^13,20,21^.

We recently employed a genome-scale analysis of codon usage to show that virulence genes of the major pathogen *Salmonella enterica* preferentially use codons that are rare in highly expressed ribosomal-protein genes, supporting efficient virulence-gene expression during nutrient limitation^22^. These virulence-associated codons include Arg-AGA/AGG, Leu-TTA/TTG, and Ile-ATA that depend on wobble-decoding, suggesting a requirement for specific tRNA modifications.

Here, we constructed a comprehensive bacterial codon–tRNA modification map to predict which modifications might preferentially support *Salmonella* virulence. We systematically tested these predictions using a series of strains with inactivating point mutations in tRNA-modifying enzymes and determined their fitness during systemic infection. Using synonymous recoded reporters, quantitative proteomics and endogenous *ssrB* recoding, we confirmed inefficient translation of specific codons as the cause of the observed virulence phenotypes. Finally, we extended our framework across bacterial pathogens and observed similar codon usage-related requirements for a small set of predictable tRNA modifications for virulence. Thus, specific tRNA modifications may provide common anti-virulence targets to control infections caused by diverse bacterial pathogens.

## Results

### Codon usage predicts required tRNA modifications for virulence genes

We previously showed that the unique codon usage of *Salmonella* virulence genes alleviates translational competition with highly expressed genes. This enables robust virulence expression even under nutrient starvation^22^. Because this program is enriched in wobble-decoded and rare codons, it may create dependence on anticodon stem-loop (ASL) tRNA modifications^1,5,6,23,24^.

To test this idea, we constructed a decoding map connecting individual codons in genes of *Salmonella enterica* serovar Typhimurium SL1344 (hereafter *Salmonella*) with respective tRNAs and their ASL modifications, based on MODOMICS^25^, current reviews^7^ and pathway-specific primary literature (Extended Data Table 1,2). The underlying decoding table is a broadly usable community tool provided as a machine-readable table together with exhaustive literature evidence (Supplementary Data 1). We then projected codon enrichment in virulence genes (Fig. 1a, Supplementary Data 2). We also quantified these relationships for individual genes by integrating the curated decoding map into CodonPipe, our genome-scale codon-usage analysis pipeline^22^. The results revealed that virulence genes were indeed enriched compared to ribosomal genes in codons that are linked to specific tRNA modifications: Cm_32_/Um_32_, Cm_34,_ k^2^C_34_, c^5^-U_34_ modification, Q_34_, and (ms^2^)i^6^A_37_ (Fig. 1b; Extended Data Fig. 1a,c,e,g,i; Supplementary Data 3,4). Thus, *Salmonella* virulence might depend on some or all these modifications.

**Fig. 1.**
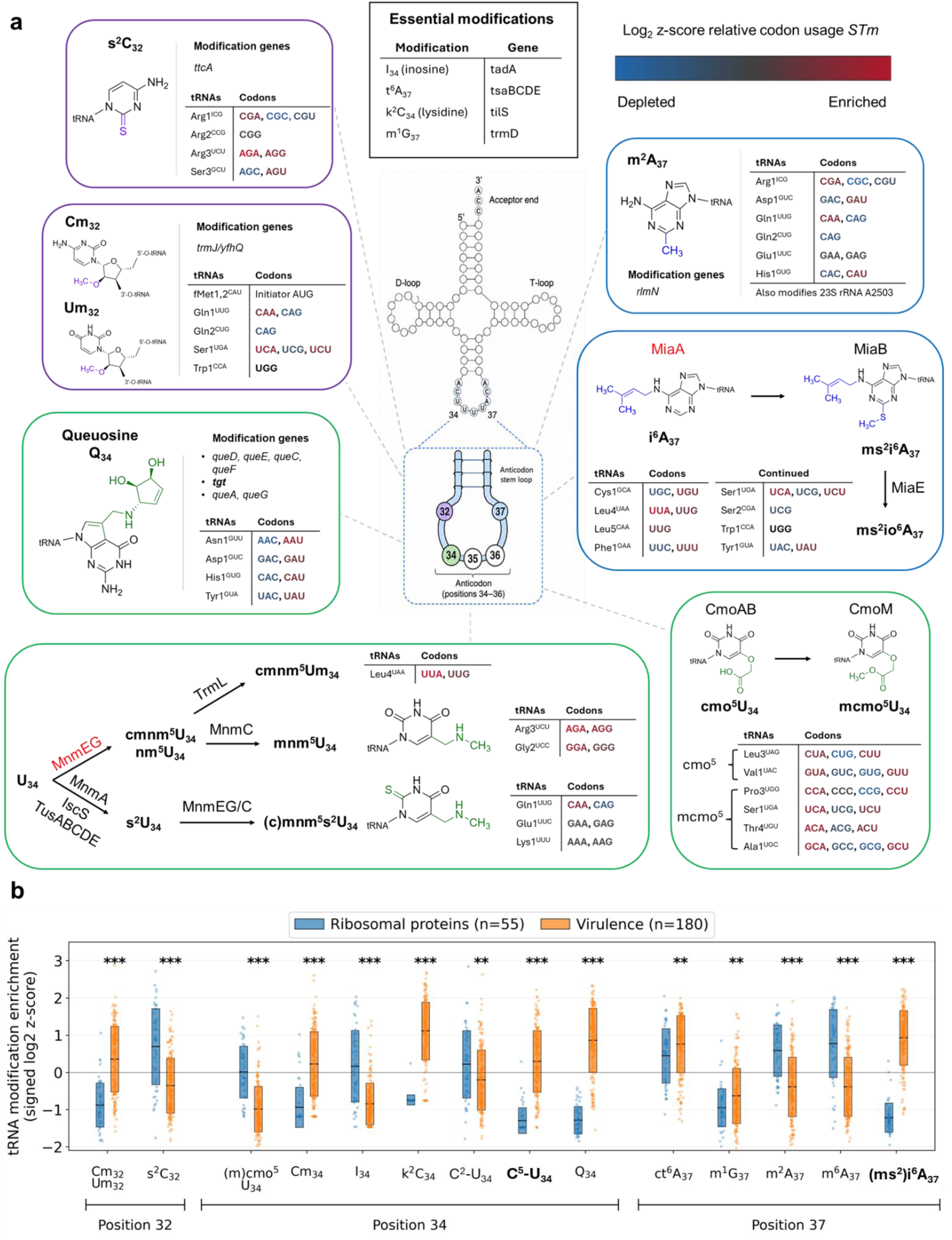
*Salmonella* virulence genes display a distinctive anticodon-loop tRNA modification signature. **a**, Curated map of bacterial tRNA modifications at anticodon-loop positions 32, 34 and 37. For each modification, the panel lists the biosynthetic enzymes, the reported tRNA substrates and the codons decoded by the modified tRNA. Codons shown in parentheses are assigned through wobble or expanded decoding. Border color denotes the modified position (purple, 32; green, 34; blue, 37). At U_34_, MnmEG-dependent C^5^-U_34_ modification and MnmA-dependent C^2^-U_34_ thiolation are treated as distinct pathways, both chemical modifications occurring on the same mature tRNA. Codon colors indicate codon-usage enrichment in *S*Tm virulence genes, expressed as signed log₂-transformed genome-normalized z-scores (scale bar). Enzyme names in red highlight modification pathways for which catalytic inactivation selectively impaired virulence (Fig. 2c). The inset box lists anticodon-loop modifications that are essential in Enterobacterales and were therefore not screened. Codon and gene assignments were curated from MODOMICS^25^, reviews^7^ and pathway-specific primary literature (Extended Data Table 1). **b**, Distribution of tRNA modification enrichment scores in *S*Tm ribosomal-protein (blue, n = 55) and virulence (orange, n = 180) genes. Each dot is one gene, boxes show median and interquartile range. Scores are the relative usage of codons decoded by tRNAs carrying the indicated modification within the relevant amino-acid families, normalized genome-wide as z-scores and transformed as sign(z) × log2(|z| + 1). Values are from the permissive assignment model, the corresponding conservative model is shown in Extended Data Fig. 1b. Statistical significance was assessed using two-sided Mann–Whitney *U*-tests with Benjamini–Hochberg correction across all modification comparisons within the analysis; **q < 0.01, ***q < 0.001. Virulence codon enrichment scores (a) and tRNA modification scores (b) are provided in Supplementary Data 2 and 3, respectively. Gene cluster definitions are provided in Supplementary Data 4.

### i^6^A_37_ and c^5^-U_34_ modifications are essential for *Salmonella* fitness in vivo

To test this idea, we deleted 19 non-essential tRNA modification enzymes spanning nearly all ASL modification pathways in *S*Tm SL1344 (Fig. 2a,b, Extended Data Table 3). We did not investigate TadA (I_34_), TsaBCDE (t^6^A_37_), TilS (k^2^C_34_) and TrmD (m^1^G_37_) that are essential in *Salmonella*^26^ (Fig. 1a). YfiC (m^6^A_37_) was excluded because its target codons are depleted in virulence genes (Fig. 1b), and RlmN because it modifies 23S rRNA in addition to tRNAs, making assignment of phenotypes to decoding difficult. Importantly, at U_34_, we distinguish MnmEG-dependent C^5^ modification from MnmA-dependent C^2^ thiolation. These chemically distinct modifications can coexist on the same mature tRNA and are further referred to as C^5^-U_34_ modification and C^2^-U_34_ thiolation (Extended Data Table 2).

**Fig. 2.**
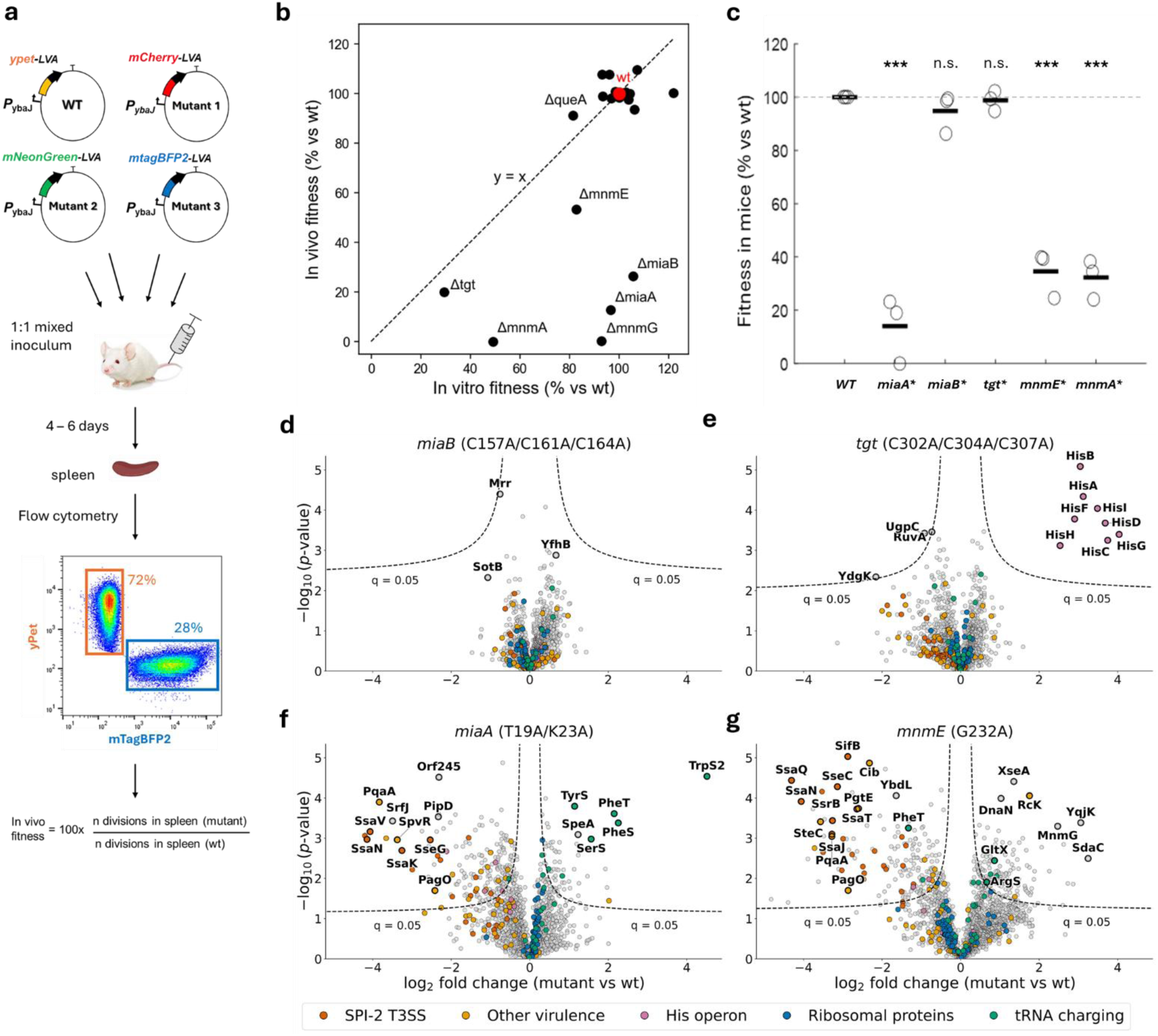
MiaA-dependent i^6^A_37_ and MnmEG-dependent C5-U_34_ modifications selectively support *Salmonella* virulence. **a**, Competitive infection scheme. Mice were co-infected with WT and up to three isogenic mutants carrying distinct constitutive fluorescent reporters. Spleens were collected after 4 days in BALB/c or 6 days in C57BL/6J *Slc11A1^r/r^* mice. Total bacterial loads were quantified by plating and relative strain abundances by flow cytometry. *In vivo* fitness was calculated from bacterial divisions during infection relative to WT. **b**, Screen of 19 tRNA modification deletion mutants for *in vitro* and *in vivo* fitness. Each point represents one mutant and shows mean fitness from three biological replicates *in vitro* and at least two BALB/c mice *in vivo*. Full values, SDs and sample sizes are provided in Extended Data Table 3. **c**, Catalytic validation of selected screen hits in the histidine-prototrophic SL1344 *hisG^L69^* background using C57BL/6J *Slc11A1^r/r^* mice. Circles represent individual mice (n = 3), horizontal bars indicate means and the dashed line indicates WT fitness (100%). Two-tailed one-sample t-tests against 100% (***P < 0.001; n.s., not significant). **d–g**, Quantitative proteomic comparisons of *miaB\** (**d**), *tgt\** (**e**), *miaA\** (**f**) and *mnmE\** (**g**) with the isogenic WT. Each dot is one protein. Selected SPI-2/T3SS-2, virulence, histidine-biosynthesis, ribosomal and tRNA-charging proteins are highlighted as indicated. Black dashed curves denote the permutation-based FDR threshold (q = 0.05). Proteomes were obtained from three independent chemostat cultures under SPI-2-inducing conditions. *miaA\**, *miaB\**, *mnmE\** and *tgt\** are the catalytic inactive mutants. Proteome data and statistics are provided in Supplementary Data 5.

We tested the 19 deletion mutants initially in competitive systemic infections in genetically susceptible BALB/c carrying dysfunctional alleles of the major resistance gene *Slc11A1* (Fig. 2a). In vivo fitness was calculated based on competitive indices in spleen 4d post-infection^27^ to enable comparison to in vitro fitness in liquid cultures in chemically defined MES-ch medium. Most mutants had limited phenotypes in vivo and in vitro indicating weak effects on decoding (Fig. 2b). However, *ΔmnmG* and *ΔmnmA* did not grow in spleen, while *ΔmiaA*, *Δtgt*, *ΔmiaB* and *ΔmnmE* were severely attenuated. *ΔmnmA* and *Δtgt* also showed substantial growth defects in vitro indicating impact beyond virulence genes. These data supported a potential virulence role for c^5^-U_34_ (MnmG), Q_34_ (Tgt), and (ms^2^)i^6^A_37_ (MiaAB) modifications, while Cm_32_/Um_32_ (TrmJ) and Cm_34_ (TrmL) had limited impact consistent with only minor decoding defects in their absence in *E. coli*^28–30^. Another predicted virulence gene-associated modification (k^2^C_34_) could not be tested because of its essentiality in vitro. In addition, C^2^-U_34_ thiolation (MnmA) showed pleiotropic effects in vitro and in vivo, consistent with our prediction (similar enrichment scores for virulence factors and ribosomal proteins). Importantly, non-predicted tRNA modifications were not required for *Salmonella* fitness during infection.

To distinguish phenotypes involving defects in tRNA modification from non-catalytic roles of these enzymes or polar effects of the genetic perturbations, we generated inactivating point mutations in *miaA*, *miaB*, *mnmA*, *mnmE*, and *tgt* (Extended Data Table 4). To test the mutants under more physiologically relevant conditions, we introduced these inactivating point mutations in histidine-prototrophic SL1344 *hisG^L69 27^* and determined in vivo phenotypes in genetically more resistant C57BL/6J *Slc11A1^r/r^* mice carrying functional *Slc11A1* alleles that restrict *Salmonella* replication through magnesium deprivation^31^. Under these conditions, functional MiaA and MnmE were specifically required for in vivo fitness (Fig. 2c). Active Tgt or MiaB were not required, while MnmA was required both in vivo and in vitro. Thus, i^6^A_37_ (MiaA) and c^5^-U_34_ (MnmE) modifications were specifically required for systemic virulence of *Salmonella*, consistent with our predictions, while Q_34_ (Tgt) and ms^2^i^6^A_37_ (MiaB) had limited impact on *Salmonella* virulence and C^2^-U_34_ thiolation (MnmA) had a pleiotropic impact. As noted above, the additional predicted k^2^C_34_ (TilS) could not be tested because it is essential in vitro^26^.

### MiaA and MnmE are required for normal SPI-2 expression

Both MiaA^11^ and MnmE (TrmE)^12^ have previously been reported as essential for *Salmonella* virulence, but the underlying mechanisms and why only these two modifying enzymes are specifically required remained unknown. To address this issue, we determined the abundance of ∼2,000 proteins in mutants *miaA* T19A K23A (*miaA*\*), *miaB* C157A C161A C164A *(miaB*\*), *mnmE* G232A (*mnmE*)*, and *tgt* C302A C304A C307A (*tgt*\*) and isogenic wild-type during growth in chemostats under infection-mimicking conditions^32^ (Fig. 2d-g; Supplementary Data 5).

*miaB\** exhibited no detectable impact on protein abundance, consistent with its wild-type level of fitness in vivo (Fig. 2d). This suggests that under-modified i^6^A_37_ was sufficient for decoding in agreement with previously observed mild phenotypes of *miaB* mutants^33^.

*tgt\** showed few proteome alterations with eight upregulated proteins encoded by the histidine-biosynthesis operon *his*GDCBHAFIE (Fig. 2e). The *his* operon leader-peptide gene *hisL* contains four CAU codons requiring the Tgt-mediated Q_34_ modification for efficient decoding^3,34^ (Fig. 1a), explaining reduced transcriptional attenuation of the operon, resulting in elevated protein levels. HisF and HisH can cause cell-division defects and filamentation^35,36^, and such effects are likely more pronounced in histidine-auxotrophic SL1344 compared to prototrophic SL1344 *hisG^L69^*, which could explain the marked in vivo phenotype of *tgt* mutants in SL1344 but not in SL1344 *hisG^L69^*. Other proteins, including histidine-regulated transporter HisJMPQ were not significantly affected. Individual virulence genes were all non-significant, but 27 of 30 quantified proteins associated with the *Salmonella* pathogenicity island 2 (SPI-2) were less abundant in the mutant (median ∼1.5-fold decrease; group-wise *q* = 3 × 10^−5^; Extended Data Table 5). These data confirmed our prediction that the Tgt-dependent tRNA modification Q_34_ supports decoding of virulence genes, but the effects were too small to result in a detectable fitness phenotype in vivo.

In contrast, *miaA*\* and *mnmE*\* mutants exhibited major proteome alterations (Fig. 2f,g), which correlated closely with each other (Pearson r = 0.67, *P* = 2 × 10^−59^). This included comprehensive suppression of SPI-2-associated proteins in both mutants: while we detected 34 SPI-2 proteins in wild-type *Salmonella*, ten such proteins became undetectable in both mutants and 23 (*miaA\**) or 24 (*mnmE\**) out of the remaining 24 SPI-2 proteins were less abundant than in the wild-type (Fig. 2f,g; Extended Data Table 5). The SPI-2 SsrA/SsrB two-component system was also strongly depleted (SsrA, ∼8-fold in both mutants; SsrB, ∼4-fold in *miaA\** and ∼10-fold in *mnmE\**), which may have contributed to the comprehensive suppression of their regulon (see below). In addition to the commonly diminished virulence factors, the two mutants showed distinct patterns among upregulated proteins. *miaA\** upregulated aminoacyl-tRNA synthetases (TrpS2, PheS, PheT, SerS, and TyrS) charging four of the six amino acids decoded by MiaA-modified i^6^A_37_ tRNAs, possibly due to relaxed transcriptional attenuation (the leader-peptide gene *pheM* upstream of *pheST* contains five Phe-TTT/TTC and one Tyr-TAC codons requiring i^6^A_37_ modified tRNAs for efficient decoding; the leader-peptide upstream of *trpS2* contains a tandem pair of Trp-TGG codons). This is consistent with recent findings in *P. aeruginosa*, demonstrating that loss of MiaA impaired decoding of Trp-UGG codons in the *trp* leader peptide, relieving repression of Trp biosynthesis^15^. *mnmE\** upregulated two aminoacyl-tRNA synthetases (GltX, ArgS) charging two amino acids decoded by MnmEG-modified tRNAs. Moreover, MnmG was more abundant, suggesting compensatory regulation at low activity of the MnmEG complex.

Thus, proteomics supported a specific requirement of i^6^A_37_ and c^5^-U_34_ modifications for efficient decoding of *Salmonella* virulence genes, whereas Q_34_ had limited effects on virulence in a histidine-prototrophic strain.

### Three signature codons correlate with the impact of tRNA modifications

Proteins with diminished abundance in the *miaA* and *mnmE* mutants formed a tight cluster in the *Salmonella* codon-usage landscape^22^ (Extended Data Fig. 2), indicating a shared codon signature.

To test the idea that this specific codon usage was indeed linked to dependency on i^6^A_37_ and c^5^-U_34_ modifications, we compared expression of synthetic *gfp* variants with codon usage approximating those of virulence genes (*gfp-VIR*) or ribosomal-protein genes (*gfp-RIB*; both variants share identical first 25 codons to minimize differences in translation initiation and transcript stability^22,37–39^; Fig. 3a). *gfp-RIB* expression was similar in wild-type *Salmonella* and in the *ΔmiaA* and *ΔmnmG* mutants, whereas *gfp-VIR* expression was suppressed in both mutants relative to wild-type (Fig. 3b,c). These data show that codon usage alone was sufficient to confer virulence-like requirements for tRNA modifications, independently of virulence regulation networks or individual virulence factors.

**Fig. 3.**
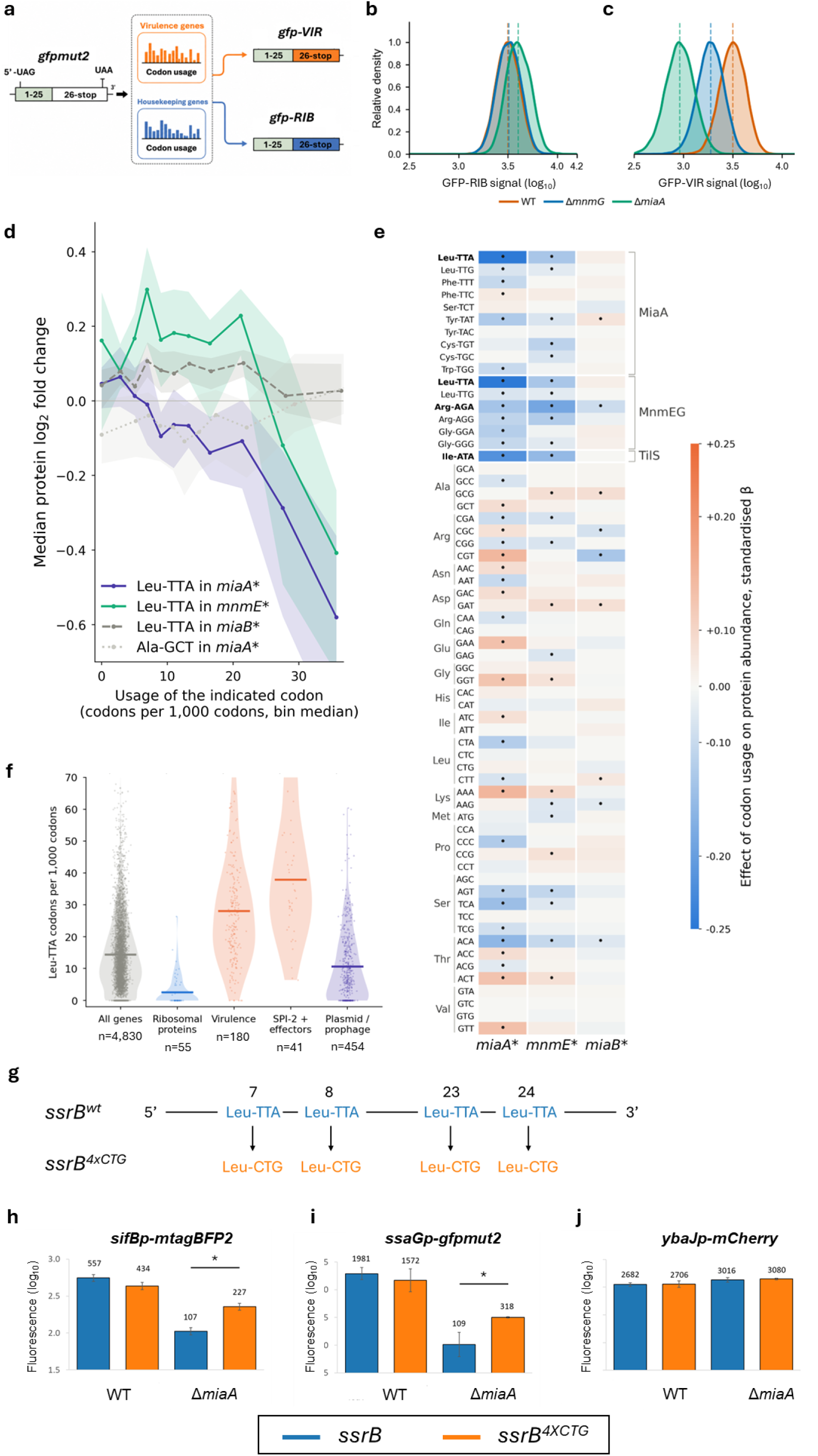
Virulence-associated synonymous codons confer tRNA modification dependence on SPI-2 expression. **a,** Synonymous *gfpmut2* reporter recoded to match the average codon usage of *S*Tm virulence (*gfpmut2-VIR*) or ribosomal-protein (*gfpmut2-RIB*) genes while retaining identical first 25 codons^22^. **b**,**c**, Flow-cytometry fluorescence distributions of *gfpmut2-RIB* (**b**) and *gfpmut2-VIR* (**c**) in the *WT*, *ΔmiaA* and *ΔmnmG* strains grown in M9 medium. **d**, Protein depletion scales with Leu-TTA usage in *miaA\** and *mnmE\** proteomes. Genes were binned by Leu-TTA (or Ala-GCT) frequency among synonymous codons. Points show median protein log_2_FC and shaded areas 95% bootstrap confidence intervals. Leu-TTA effects are shown for *miaA\**, *mnmE\** and *miaB\** proteomes. Ala-GCT in *miaA\** serves as a codon-specificity control. **e**, Codon associations with protein abundance in *miaA\**, *mnmE\** and *miaB\** proteomes. All 61 sense codons were tested by linear regression adjusting for GC3 (G+C at third position) content and gene length. Note that Leu-TTA/TTG appear under both MiaA and MnmEG because the decoding tRNA carries both modifications, values are identical. Displayed codons are grouped by associated tRNA modification pathway as indicated, then listed alphabetically. Colors indicate standardized regression coefficients β (blue, depletion; orange, enrichment) and dots indicate q < 0.05 after Benjamini–Hochberg correction across 244 tests. Codon associations are shown for *miaA\**, *mnmE\** and *miaB\** and the full dataset is provided in Supplementary Data 6. **f**, Distribution of Leu-TTA usage across *S*Tm gene clusters. Dots represent coding sequences; violins show distributions and horizontal bars indicate medians. Functional classes were compared with the genome-wide distribution by two-sided Mann–Whitney U-tests (all P < 10^−14^). n = 4,830 of the 4,868 annotated coding sequences; 38 sequences shorter than 30 sense codons were excluded. **g**, Synonymous recoding scheme of Leu-TTA tandem codon pairs in *ssrB*. Four Leu-TTA codons at positions 7, 8, 23 and 24 were replaced by the synonymous Leu-CTG codon, leaving the SsrB protein sequence unchanged. **h**–**j**, Effect of synonymous *ssrB* recoding on SPI-2 expression. SPI-2 expression was tested in MES-ch-inducing medium in the *WT* and *ΔmiaA* mutant using chromosomal *sifBp-mtagBFP2* (**h**), plasmid-borne *ssaGp-gfpmut2* (**i**) and a constitutive control *ybaJp-mCherry* (**j**) reporters. Bars show log_10_-transformed mean fluorescence from two independent biological replicates ± s.d. Statistical significance was assessed by two-tailed Student’s t-tests. FC, fold-change; *miaA\**, *miaB\**, *mnmE\** and *tgt\** are the catalytic inactive mutants. Gene cluster definitions (f) are provided in Supplementary Data 4. Codon-level statistics (d,e) are provided in Supplementary Data 6.

To determine which codons conferred dependence on i^6^A_37_ and C^5^-U_34_ modifications, we related the frequency of each codon within a gene to the fold suppression of the corresponding protein in the mutants. For instance, genes with high frequency of codon Leu-TTA were strongly suppressed in both *miaA\** and *mnmE\** but not *miaB*\*, whereas Ala-GCT frequency had no impact on protein abundance alterations in these mutants (Fig. 3d). A comprehensive analysis of all codons revealed associations predominantly among codons linked to i^6^A_37_ and c^5^-U_34_ modifications consistent with our predictions (Fig. 3e, Supplementary Data 6). The strongest associations involved Leu-TTA (i^6^A_37_ and c^5^-U_34_ modifications) and Arg-AGA (c^5^-U_34_ modification). A key role of Leu-TTA was consistent with its differential abundance (high in virulence genes, low in ribosomal genes; Fig. 3f). Other codons linked to these modifications had small or non-detectable associations indicating that their decoding was less affected by under-modified tRNAs. Varying impact of tRNA modifications on decoding of different codons has been previously observed, including a dominant impact of Leu-TTA/TTG on MiaA-dependent translation of *rpoS* in *E. coli*^9,40^, and of Arg-AGA on MnmEG-dependent expression of *pINV* virulence genes in *Shigella*^10^ (Extended Data Table 6).

Unexpectedly, Ile-ATA-rich genes were also depleted in both *miaA\** and *mnmE\**, although the linked tRNA modification k^2^C_34_ should not be affected in these mutants. A likely explanation is the co-enrichment of the three key codons Leu-TTA, Arg-AGA, and Ile-ATA as part of the codon-usage signature of virulence genes (Fig. 4c; Extended Data Fig. 3c,d)^22^. Thus, Leu-TTA and Arg-AGA may directly confer sensitivity to loss of i^6^A_37_ or C^5^-U_34_ modifications, whereas Ile-ATA is a co-enriched bystander with no independent effect.

**Fig. 4.**
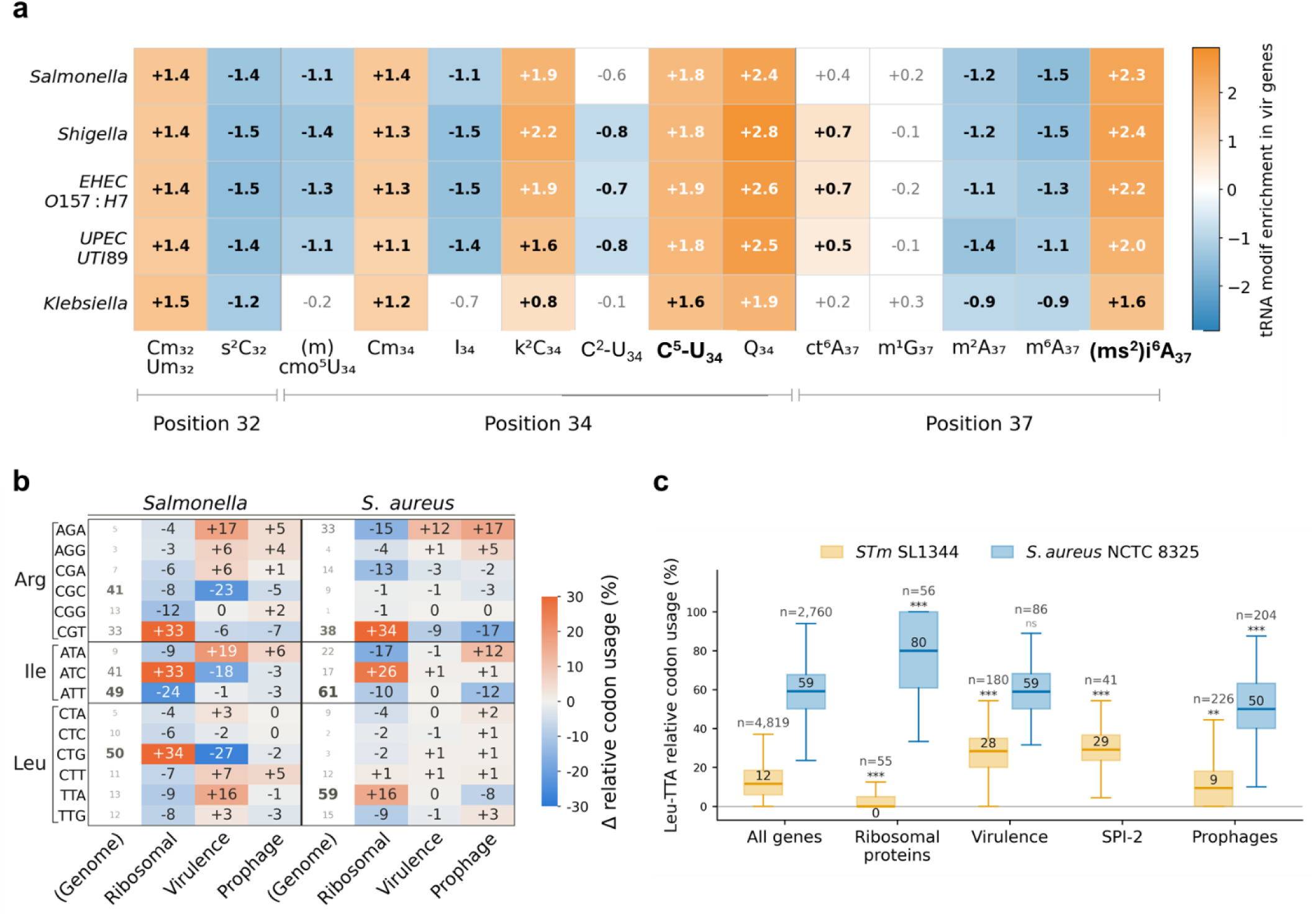
Virulence-associated tRNA modification signatures are conserved across bacterial pathogens. **a**, Difference in tRNA modification enrichment between virulence and ribosomal-protein genes across five Enterobacterales pathogens (virulence minus ribosomal-protein median score, permissive model). Orange indicates enrichment in virulence genes and blue enrichment in ribosomal-protein genes. Bold labels indicate the modification pathways for which catalytic inactivation selectively impaired virulence (Fig. 2c). Bold values indicate q < 0.001 after Benjamini–Hochberg correction. **b**, Synonymous codon usage in *S*Tm SL1344 and *S. aureus* NCTC 8325. Genome values indicate relative usage within each synonymous codon family (%), with font size scaled to frequency. Ribosomal, virulence and prophage columns show differences from the genome-wide average (blue, depleted, orange, enriched). The 15 arginine, isoleucine and leucine codons are shown; the remaining 44 of the 59 synonymous codons are provided in Extended Data Fig. 3b. **c**, Gene-level Leu-TTA usage across functional classes in *S*Tm (yellow) and *S. aureus* (blue). Numbers above groups indicate gene counts and values within boxes indicate median Leu-TTA usage (%). SPI-2 is specific to *S*Tm and therefore has no corresponding *S. aureus* group. Statistical significance was assessed using two-sided Mann–Whitney U-tests; n.s., not significant, **q < 0.01, ***q < 0.001. Gene cluster definitions are provided in Supplementary Data 4.

### Two Leu-TTA tandems in *ssrB* contribute to SPI-2’s dependency on i^6^A_37_

In addition to the common codon usage of virulence genes impairing translation in the absence of i^6^A_37_ or C^5^-U_34_ modifications, low levels of the SPI-2 master regulator SsrB (see above) could reduce transcription of the SPI-2 regulon. To test this idea, we utilized two transcriptional fusions to canonical SsrB-dependent promoters (chromosomal *sifBp-mtagBFP2*, plasmid-borne *ssaGp-gfp*). Both fusions showed diminished activity in Δ*miaA* as compared to wild-type *Salmonella*, while a constitutively active *ybaJp-mCherry* fusion remained unaffected (Fig. 3h–j). This confirmed a significant contribution of reduced SsrB activity on diminished SPI-2 regulon expression.

*ssrB* contains two Leu-TTA tandem pairs at codons 7–8 and 23–24. Because of the key role of Leu-TTA and enhanced impact of tandem modification-sensitive codons^40,41^, we exchanged these four Leu-TTA codons to Leu-CTG (*i.e.*, without changing the amino acid sequence; Fig. 3g). This *ssrB* recoding partially rescued the diminished activities of *sifBp* and *ssaGp* expression, while the constitutive *ybaJp* remained unchanged (Fig. 3h–j). Thus, four synonymous codons in a master regulator contributed significantly to the dependence of an entire virulence program on i^6^A_37_. However, *ssrB* recoding was insufficient to restore in vivo fitness of the *miaA* mutant (competitive indices: *miaA\**, 1.7×10^−5^ ± 1.1×10^−5^; *miaA*\* *ssrB*^4xCTG^ 4.0×10^−5^ ± 3.2×10^−5^; *P* = 0.21, n = 3). This may be related to both incomplete rescue of SPI-2 transcription and still inefficient translation of individual Leu-TTA-rich transcripts.

### Virulence-associated tRNA modification signatures are conserved across *Enterobacterales* pathogens

*miaA*, *mnmA*, *mnmE*, *mnmG*, *tilS* and *tgt* are broadly conserved across bacterial phyla (Extended Data Fig. 3a). To determine if they play similar roles in the virulence of other pathogens (and may thus qualify as broader anti-virulence targets), we applied our comprehensive in silico analysis of tRNA modifications to *Shigella flexneri 2a*, enterohemorrhagic *E. coli* O157:H7 Sakai, uropathogenic *E. coli* UTI89, and *Klebsiella pneumoniae* HS11286. Despite substantial differences in genome content and virulence repertoires, all five pathogens displayed common separations between virulence and ribosomal-protein genes (Fig. 4a; Extended Data Fig. 1; Supplementary Data 4). As in *Salmonella*, virulence genes were consistently enriched in codons associated with i^6^A_37_, Q_34_, c^5^-U_34_ and k^2^C_34_ modifications.

This signature is consistent with reported virulence phenotypes of MiaA or MnmEG mutants in *S. flexneri*^10^ and *S.* Typhimurium^8,11,12^, and might also extend to extraintestinal pathogenic *E. coli*^13^, *P. aeruginosa*^14^, and *S. pyogenes*^42^ (Extended Data Table 6). Together, these observations suggest overlapping virulence-associated codon usage and corresponding tRNA modification requirements across diverse major bacterial pathogens.

However, we also identified an exception, the AT-rich (67.1%) Gram-positive pathogen *Staphylococcus aureus*. Synonymous codon usage differed markedly compared to *Salmonella* (Fig. 4b; Extended Data Fig. 3b). As an example, Leu-TTA is relatively rare in *Salmonella* and nearly excluded from ribosomal-protein genes yet enriched in virulence genes. In *S. aureus*, by contrast, it accounts for 59% of leucine codons genome-wide and 75% in ribosomal-protein genes, while showing no enrichment in virulence genes. This opposite codon usage was not a generic consequence of *S. aureus*’s high AT content, because Arg-AGA and Ile-ATA retained preferential enrichment in virulence and prophage genes relative to ribosomal-protein genes (although both codons had high overall frequencies; Extended Data Fig. 3c,d). *S. aureus* virulence would therefore be predicted to be less dependent on i^6^A_37_, which is confirmed by mild *miaA*-phenotypes in various mouse infection models^43,44^.

Conversely, the high basal frequencies of Arg-AGA and Ile-ATA predict broad requirements for MnmEG-dependent C^5^-U_34_ modification and TilS-dependent k^2^C_34_, respectively. Consistent with this, *mnmG* (*gidA*), *mnmE* (*trmE*) and the tRNA-Ile modification pathway have been identified as essential for *S. aureus* in vitro^45^. Together these observations demonstrate a clear link between virulence-gene codon usage and tRNA-modification dependence, in which the identity of the critical codons is set by each species’ codon architecture (Fig. 5). These links can be predicted by our in silico pipeline for any sequenced pathogen genome (https://doi.org/10.5281/zenodo.21303183).

**Fig. 5.**
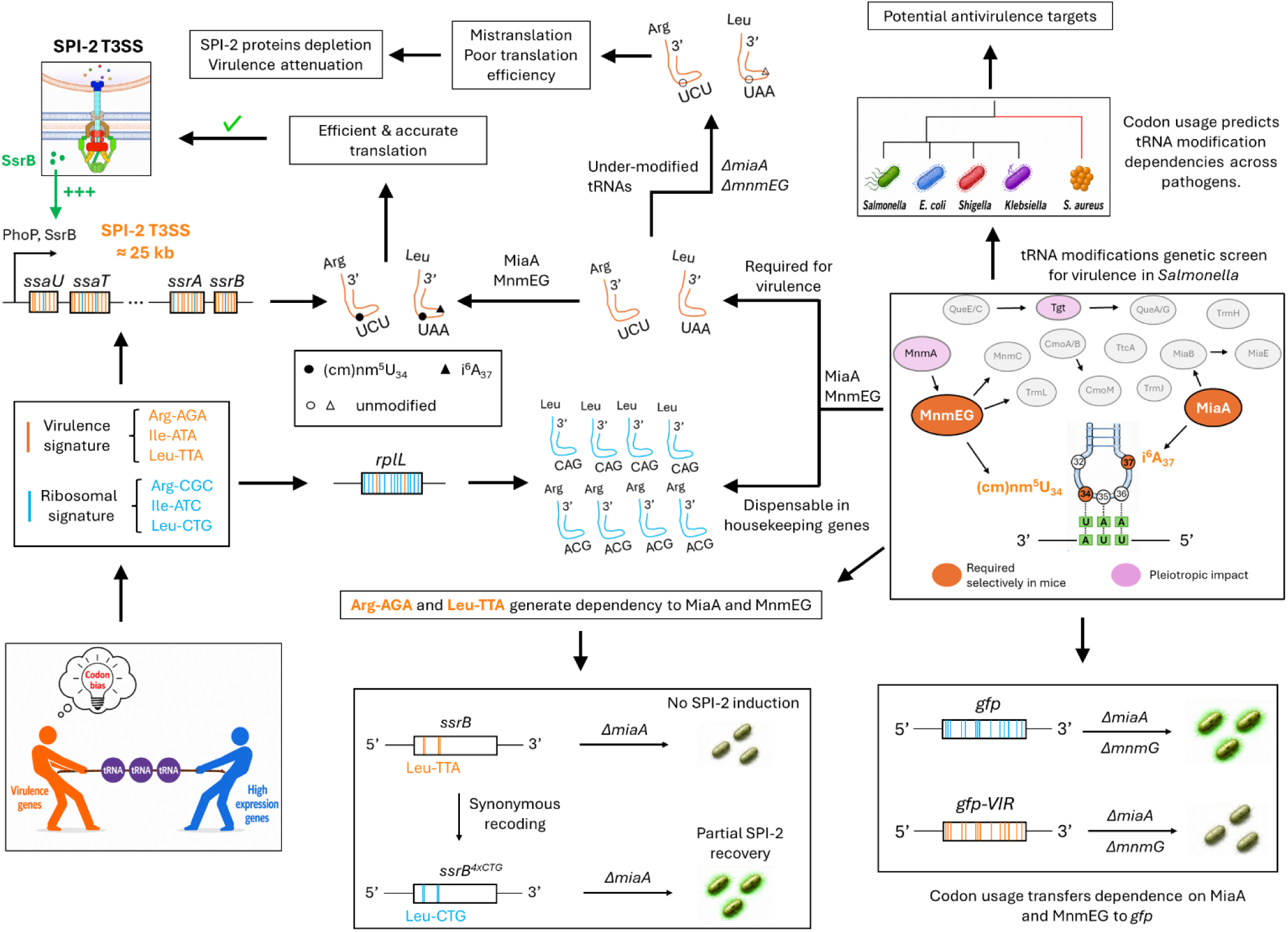
Model linking virulence-associated codon usage to tRNA-modification dependence. *Salmonella* virulence genes preferentially use synonymous codons that minimize competition with highly expressed genes but increase dependence on specific anticodon-loop tRNA modifications. MiaA-dependent i^6^A_37_ and MnmEG-dependent C^5^-U_34_ modifications support efficient decoding of modification-sensitive codons, in particular Leu-TTA and Arg-AGA, thereby sustaining expression of virulence proteins and synthesis of the SPI-2 T3SS. Reduced translation of the Leu-TTA-rich SPI-2 regulator SsrB further amplifies these effects through transcriptional suppression of the SPI-2 regulon. Synonymous recoding can alter this dependence without changing protein sequence. Across bacterial pathogens, the general relationship between virulence-associated codon usage and tRNA modification is conserved, whereas the identity of individual codons can vary with species-specific codon usage.

## Discussion

tRNA modifications are necessary for efficient translation of various codons. Some of these tRNA modifications are essential for virulence of diverse pathogens in various in vitro and in vivo infection models^8,10–14,17^. Why particular pathways are required for virulence, and whether these dependencies can be predicted from virulence-gene codon usage, has remained largely unclear. Here, we show that the distinctive synonymous codon architecture of *Salmonella* virulence genes couples their expression to a few specific tRNA modifications.

Based on a codon-tRNA modification map, we predicted a maximum of six modifications to be specifically associated with virulence. However, some of these modifications are known to have limited impact on decoding. This was confirmed by testing 19 mutants affecting non-essential anticodon-loop modification pathways in infected mice, with most mutants showing wild-type levels of fitness. However, some mutants showed strong impact: MiaA-dependent formation of i^6^A_37_ was required, whereas downstream conversion to ms^2^i^6^A_37_ by MiaB was dispensable consistent with its limited impact on decoding. Likewise, MnmEG-dependent C^5^-U_34_ modification was required in mice, whereas downstream MnmC processing was dispensable. These findings confirm and extend recent observations that MnmEG supports expression of genes encoded on the *Shigella* pINV virulence plasmid^10^. Cm_32_/Um_32_ and Cm_34_ were not required, consistent with their generally limited impact on decoding. Q_34_ had a detectable minor impact on decoding of virulence genes but this was dispensable for in vivo fitness. Thus, virulence depended only on a small set of tRNA modifications in line with our genome-based predictions.

*Salmonella* virulence depended on i^6^A_37_ and C^5^-U_34_ modification through two related mechanisms. First, these tRNA modifications supported translation of genes sharing the codon signature of virulence genes. Second, they also supported transcription of SPI-2 genes by increasing the abundance of the master regulator SsrB, which contains two tandem pairs of modification-sensitive codons. In addition, i^6^A_37_ and Q_34_ are required to maintain functional regulation of attenuation-controlled operons encoding aminoacyl-tRNA synthetases (*pheST*, *trpS2*) and histidine biosynthesis (*his*), consistent with recent work in *P. aeruginosa* revealing that the absence of i^6^A_37_ modification disrupted *trp* attenuation, in turn causing broadly altered virulence-gene expression^15^.

This raises the question of why virulence genes are enriched for codons that depend on these specific modifications. Our previous work showed that *Salmonella* virulence genes avoid synonymous codons heavily used by highly expressed housekeeping genes, reducing competition for charged tRNAs and sustaining virulence-gene translation during nutrient limitation^22^. This results in enrichment of rare and/or wobble codons which are uncommon in ribosomal-protein genes. Such codons can require ASL modifications that stabilize codon-anticodon interactions, support efficient decoding, and enable wobble pairing. Dependence on MiaA and MnmEG may therefore represent a consequence of a codon signature selected primarily to maintain robust virulence-gene expression under host-imposed nutrient limitation. In this view, tRNA modification dependence may not be adaptive in itself but rather emerge from the complementary codon usage of highly expressed housekeeping and virulence genes. We extended our genome-wide analysis across Enterobacterales and observed shared predicted tRNA-modification dependencies, including i^6^A_37_ and C^5^-U_34_ modifications. Previous studies linking MiaA or MnmEG to pathogenicity in several species support broader relevance of these pathways. A notable exception is *Staphylococcus aureus* with opposite patterns and tRNA-modification phenotypes. Thus, species-specific codon usage can reassign individual codons between functional gene clusters, making tRNA modification dependence an emergent property of each species’ codon architecture rather than a fixed feature of the modification itself. Our machine-readable codon-tRNA-modification map and open-access pipeline therefore provide a broadly usable framework to predict modification dependencies directly from genome sequence. Conversely, genes sharing modification-sensitive codon signatures could be identified as candidate virulence genes or other functional modules with marked modification requirements.

The shared dependency of virulence on MiaA and MnmEG across pathogens might enable anti-virulence strategies for infection control. However, humans possess homologous enzymes TRIT1 and the mitochondrial GTPBP3–MTO1 system that share reaction mechanisms and active-center structures^46–49^. Nevertheless, selective targeting of bacterial enzymes may be possible, similar to selective inhibition of bacterial but not human dihydrofolate reductases by the clinically successful antibiotic trimethoprim^50^.

Together, our results establish synonymous codon architecture as a mechanistic link between bacterial virulence and tRNA modification. This dependence is chemically specific, distributed across many virulence proteins, amplified by global regulators such as SsrB, and reshaped by species-specific codon usage (Fig. 5).

## Methods

### Bacterial strains and culture conditions

All strains were derived from *Salmonella enterica* serovar Typhimurium SL1344 (further referred to as *S*Tm)^51^ or from SME51^27^, a prototrophic *hisG^Leu69^* derivative of SL1344. Strains and plasmids used in this study are listed in Supplementary Data 7. Bacteria were grown in Lennox lysogeny broth (LB), M9 medium or MES-ch medium. M9 medium contained 1× M9 salts, 0.4% glucose and 2 mM MgSO4, and was supplemented with 0.05% casamino acids. MES-ch is a SPI-2 inducing medium prepared as described previously^52^ and contains 100 mM 2-(N-morpholino)ethanesulfonic acid (MES), 5 mM KCl, 15 mM NH4Cl, 0.5 mM K2SO4, 1 mM KH2PO4, 50 μM MgSO4, 0.02% casamino acids, 0.02% glycerol, 0.0042% N-acetyl-glucosamine, 0.003% glucose, 0.0018% glucosamine, 0.005% histidine and 25 mM NaHCO3, adjusted to pH 5.5.

When indicated, antibiotics were added at 50 µg/mL kanamycin or 10 µg/mL chloramphenicol to maintain plasmid selection. Histidine was added at a final concentration of 0.005% when culturing *S*Tm SL1344, which is auxotrophic for histidine. For proteomic analyses under infection-mimicking conditions, *Salmonella* was grown in MES-ch in minichemostats^53^ at a controlled generation time of 6 h with continuous aeration using 10% O_2_ and 5% CO_2_ as described previously^54^.

### Strain and plasmid construction

The tRNA modification deletion-mutant library was generated using the mini-λ recombineering system^55^. A kanamycin-resistance cassette flanked by regions homologous to the target locus was amplified by PCR and electroporated into strains expressing λ Red recombination proteins. Recombinants were selected on kanamycin, and the mini-λ element was cured by two successive passages at 37 °C. Deletions were confirmed by sequencing. For the initial screen, each targeted tRNA modification enzyme gene was replaced entirely by the kanamycin-resistance cassette, which was retained in the resulting strain. Nineteen mutants were constructed, covering all non-essential tRNA modification pathways targeting anticodon-loop positions 32, 34 and 37 in *S*Tm, together with *ΔtrmH* (Gm18), a modification outside the anticodon loop. The complete mutant collection affected modification pathways and corresponding in vitro and in vivo fitness values are provided in Extended Data Table 3.

Catalytically inactive alleles and the recoded *ssrB* strain were generated by scarless allelic exchange. Substitutions were selected from previous biochemical and structural studies (Extended Data Table 4): *miaA* T19A K23A, combining residues implicated in prenyl-transfer catalysis and substrate binding^56,57^; *miaB* C157A C161A C164A, disrupting the conserved [4Fe–4S]-binding motif^58^; *mnmA* Q151E, impairing U_34_ recognition^59^; *mnmE* G232A, disrupting the conserved GTPase motif required for MnmE function^60^; *mnmG* G13A G15A, disrupting the Rossmann-fold FAD-binding motif^61^; and *tgt* C302A C304A C307A, disrupting the zinc-binding site required for productive tRNA interaction^62^. All recovered alleles were verified by Sanger sequencing. Despite repeated attempts, *mnmG* G13A G15A could not be recovered. Because MnmE and MnmG function cooperatively and are both required for the mature C^5^-U_34_ modification^46,63^, *mnmE* G232A was used as the catalytic proxy for this pathway.

For recoding of Leu-TTA at positions 7,8,23,24 into Leu-CTG in *ssrB*, integration plasmids containing 700-bp homology regions flanking the targeted mutations were used. Mutants were obtained through two consecutive single crossover events, using kanamycin resistance for positive selection and *sacB*-mediated sucrose sensitivity for counterselection, as described previously^64^.

The previously described recoded *gfpmut2* reporter plasmids FreGo911 and FreGo914^22^ encode *gfpmut2* variants recoded to match the codon usage of either *S*Tm virulence genes (*gfpmut2-VIR*) or highly expressed ribosomal-protein genes (*gfpmut2-RIB*). The first 25 codons were identically optimized in both constructs to minimize differences in translation initiation and transcript stability. Recoded gfpmut2 genes were expressed from the constitutive proD promoter^65^ on a low-copy pSC101 kanamycin-resistance plasmid. The same plasmids carried a second cassette expressing an IDT codon-optimized *mScarlet-i3* gene from the constitutive *PybaJ* promoter^66^ and served as a non-recoded internal fluorescent control.

For the chromosomal SPI-2 transcriptional reporter, the *sifB* cds was replaced by *mtagBFP2* using scarless allelic exchange as described above. *sifB* is strongly and continuously induced during infection^67^ as part of the SPI-2 regulon, and deletion of *sifB* has no significant impact on *S*Tm virulence^68^. The plasmid-based SPI-2 transcriptional reporter pOPC118 was constructed on a low copy pSC101 plasmid and contained a *PssaG-gfpmut2* transcriptional fusion carrying a C-terminal OVA degradation tag^68^, together with a constitutive PybaJ-*mcherry* cassette used as a bacterial marker.

### Mouse infection experiments

A schematic illustration of mouse infection procedure is shown in Fig. 2a. BALB/cAnNCrl or C57BL/6J *Slc11A1^r/r^* mice^31^ were co-infected at a 1:1 ratio by tail-vein injection with 100 μL PBS containing 500–5,000 CFU per strain of *S*Tm wt and isogenic mutants grown to late-log phase in Lennox LB. The inoculum size and input strain ratio were determined by plating for each infection. Male and female mice, 10-15 weeks of age, were used without distinction. Mice were scored daily for disease severity using predefined criteria covering spontaneous behavior, provoked behavior, physical appearance, clinical signs, hydration status, and grip strength.

Four days after infection for BALB/c mice, or six days after infection for C57BL/6J *Slc11A1^r/r^* mice, animals were euthanized with CO_2_ and spleens were collected. Spleens were homogenized in ice-cold PBS containing 0.2% Triton X-100 and samples were kept on ice until analysis. Large host cell fragments were removed by centrifugation at 500 × g for 5 min. The supernatant contained more than 90% of the total *Salmonella* population as dispersed single cells. Total bacterial loads were quantified by plating on LB agar, and relative ratios of co-infected strains were determined by flow cytometry, based on constitutive plasmid-encoded single-color fluorescent reporters specific to each strain.

Competitive index (CI) values were determined as the output bacterial ratio (mutant-out/wt-out) divided by the inoculum ratio (mutant-in/wt-in). Because CI values diverge with ongoing replication, and because *Salmonella* replicates more slowly in *Slc11a1^r/r^* mice than in susceptible animals^31^, CI values underestimate mutant phenotypes in C57BL/6 *Slc11A1^r/r^* mice. We therefore estimated the number of bacterial divisions and generation time in vivo from log_2_(FI), where FI is the fold increase from the estimated initial splenic colonization, assumed to be approximately 20% of the inoculum^69^, to the final organ load. The fitness of co-administered wild-type *Salmonella* was set to 1, and relative mutant fitness in vivo was calculated as the number of bacterial divisions over the course of infection normalized to that of the wild type.

All animal experiments were approved by the Kantonales Veterinäramt Basel (license 2239) and performed in accordance with local regulations and Swiss animal protection law. Mice were housed at 22°C (−2/+3°C), 55% ± 10% relative humidity, and a 12 h/12 h light/dark cycle. Animals were infected at 10–15 weeks of age. Sample size was estimated using a sequential statistical design. Effect size and variance were based on previous studies^31,52^, and final group sizes were adjusted to achieve adequate statistical power. Experiments were not randomized, and investigators were not blinded to strain identity. Strain assignment was determined by the constitutive fluorescent reporter carried by each strain and read out automatically by flow cytometry.

### Quantitative proteomic analyse

*S*Tm SME51 and isogenic mutants *miaA(T19A/K23A)*, *miaB(C157A/C161A/C164A)*, *mnmE(G232A)*, and *tgt(C302A/C304A/C307A)* were grown in chemostats in MES-ch medium at pH 5.5, with a controlled generation time of 6 h and continuous aeration with 10% O_2_ and 5% CO_2_ to mimic infection-like conditions as previously described^54^. Samples were harvested by centrifugation, washed in PBS, snap-frozen and stored at −80°C until processing.

Bacterial pellets were resuspended in lysis buffer containing 5% sodium dodecyl sulfate, 10 mM tris(2-carboxyethyl)phosphine, and 100 mM triethylammonium bicarbonate. Samples were incubated at 95°C for 10 min and sonicated for 20 cycles of 30 s on/30 s off in a Bioruptor system (Diagenode). Protein concentration was determined by tryptophan fluorescence (excitation 278 nm, emission 354 nm; Infinite M Plex, Tecan). Proteins were alkylated with 20 mM iodoacetamide for 30 min at 25°C with gentle shaking in the dark. Aliquots containing 20 µg protein were processed using S-Trap columns (ProtiFi) and digested with trypsin. Peptide concentration was determined using a UV-based assay on an Infinite M Nano instrument (Tecan).

Peptides were separated on a Dionex UltiMate 3000 nanoLC system coupled online to an Orbitrap Exploris 480 mass spectrometer (Thermo Fisher Scientific). Peptides were loaded onto 20 cm × 75 µm ID capillary columns packed in-house with 1.9 µm Reprosil-Pur C18 resin (Dr. Maisch). The column temperature was maintained at 50°C. Solvent A consisted of 0.1% formic acid in water and solvent B consisted of 0.1% formic acid in acetonitrile. Peptides were separated at a flow rate of 300 nL/min using a 60 min gradient from 2% to 12% solvent B over 5 min, then to 35% solvent B over 45 min, and to 50% solvent B over 10 min. Columns were subsequently washed with 95% solvent B and re-equilibrated in solvent A. The spray voltage was set to 2.5 kV, funnel RF to 40, and capillary temperature to 275°C. Data were acquired in positive-ion centroid mode using data-independent acquisition. Full MS scans were acquired at 120,000 resolution with normalized AGC target of 300%, a maximum injection time of 45 ms and a scan range of 350–1400 m/z. Fragment spectra were acquired at 15,000 resolution with a normalized AGC target of 1000%, a maximum injection time of 22 ms and normalized collision energy of 28%. DIA spectra were acquired using 63 isolation windows of 9 Da with 1 Da overlap.

Raw data were analyzed in Spectronaut v16.1.220730.53000 (Biognosys) using directDIA against a FASTA database containing the *S*Tm SL1344 proteome and common contaminants. Carbamidomethylation of cysteine was set as a fixed modification, whereas methionine oxidation and protein N-terminal methionine excision were allowed as variable modifications. Results were filtered at 1% false discovery rate at the precursor, peptide and protein levels. Downstream analyses were performed in Perseus v3.0^70^ and using Python scripts. All mass spectrometry proteomics data associated with this manuscript have been deposited to the ProteomeXchange consortium via MassIVE (https://massive.ucsd.edu) with the accession numbers MassIVE MSV000103162 / PRIDE PXD083883.

Protein abundance was calculated from protein group quantities according to the total protein approach^71^. Approximately 2,000 proteins were quantified per comparison, with proteins retained for statistical analysis when detected in at least two of three biological replicates in both the mutant and wild type. Differential abundance was assessed using two-sample t-tests on log2-transformed abundances in Perseus^70^, with s0 = 0.1 and permutation-based false-discovery-rate control at 5% using 250 randomizations. Proteins outside the permutation-based 5% FDR boundary were considered significantly changed and their q values are given in Supplementary Data 5. No individual protein in the *miaB*\* proteome crossed this threshold.

Functional-category analysis of the mutant proteomes (Extended Data Table 5). Proteins were assigned to functional categories as listed in Supplementary Data 4. For each mutant and category, Extended Data Table 5 reports the number of proteins quantified in the comparison, the median log_2_ fold-change, the number with a negative fold-change, a one-sided exact binomial sign test of that number against an expectation of one half, and a two-sided Mann–Whitney U test of the category’s fold-changes against all other proteins quantified in the same comparison. The sign test counts the direction of change rather than statistical significance, so that under a null of no category effect the number of negative changes is binomial(n, 0.5). Counting only proteins passing the FDR threshold would make the test conservative and dependent on that threshold. The Mann–Whitney comparison uses all other proteins in the same comparison rather than a fixed background, so it is unaffected by differences in proteome coverage between mutants. P values were corrected by the Benjamini– Hochberg procedure across all categories within each mutant, separately for the two tests.

### Codon-resolved analysis of proteome responses

Protein log_2_ fold-changes (mutant versus wild-type, MES-ch chemostat) were matched to SL1344 locus tags. Overall, 84% of quantified proteins mapped (1,637–1,679 per strain), and unmapped entries showed no fold-change bias. Codon counts were computed and expressed per 1,000 sense codons. For the dose–response curve (Fig. 3d), genes were ranked by the frequency of the codon of interest and cut at the 10th, 20th … 80th, 90th and 95th percentiles. The median log₂ fold-change of each bin was bootstrapped 2,000 times to obtain a 95% percentile interval, and each bin was plotted at its median codon frequency, such that the horizontal axis represents codon dose rather than rank. Ala-GCT was chosen as the codon-specificity control because none of the tRNAs decoding it carries any of the modifications disrupted in this study. For the codon matrix (Fig. 3e), because codon frequency covaries with genomic base composition, each of the 61 sense codons was modelled separately in each of the four mutant proteomes, with GC3 (G + C content at third position) and gene length included as covariates. Each model was fitted by ordinary least squares as log_2_FC ∼ β₁·codon frequency + β₂·GC3 content + β₃·log₁₀ gene length. β₁ is reported, P values are two-sided t-tests on β₁, and q values are Benjamini–Hochberg adjusted across all 244 codon × mutant tests; 95 tests were significant at q < 0.05 (Supplementary Data 6). Codons were assigned to modification pathways according to the decoding table (Extended Data Table 1; Fig. 1a).

### Flow cytometry

For competitive mice infections and fluorescence analyses of recoded *gfpmut2* and SPI-2 transcriptional reporters, fluorescence was recorded in a BD LSR Fortessa II equipped with 405-nm, 488-nm, and 561-nm lasers (Becton Dickinson), using thresholds on side scatter (SSC-H) to exclude electronic noise and gating on SSC-A vs SSC-H to exclude non-singlet events. For fluorescence acquisition, the following channels were used: mTagBFP2 (Ex 405, Em 470/20-H), GFPmut2 (Ex 488, Em 514/30-H), mNeonGreen (Ex 488, Em 512/25-H), Ypet (Ex 488, Em 542/18-H), mScarlet-i3 (Ex 561, Em 582/15-H) and mCherry (Ex 561, Em 617/73-H). Data were processed with FlowJo 10.10.1, MATLAB R2024a and Python v3.11.13.

### Calculation of tRNA modification-enrichment scores

Analyses were performed using CodonPipe, a previously described framework for genome-scale codon-usage analysis^22^. CodonPipe was extended in this study to quantify codon-usage signatures associated with anticodon-loop tRNA modifications. The updated CodonPipe platform (version 1.1) is distributed through a graphical user interface and is available on Zenodo (DOI: 10.5281/zenodo.21303183).

Codon assignments were derived from a manually curated bacterial codon–anticodon–tRNA modification table linking each sense codon to compatible tRNAs, anticodons, tRNA modifications at positions 32, 34 and 37, and the corresponding tRNA modification enzymes (Extended Data Table 1). This decoding table was curated from MODOMICS^25^, available reviews^7^ and primary literature and is summarized in Fig. 1a. Related mature modifications belonging to the same biosynthetic pathway were grouped into pathway-level classes (Extended Data Table 2), hereafter referred to as tRNA modification pathways. Cm_32_ and Um_32_ were combined as a position-32 methylation class; cmo⁵U_34_ and mcmo⁵U_34_ as (m)cmo⁵U_34_; i⁶A_37_ and ms²i⁶A_37_ as i⁶A_37_/ms²i⁶A_37_; MnmEG- and MnmC-dependent C^5^-U_34_ modifications cmnm⁵U_34_, nm⁵U_34_, mnm⁵U_34_, and cmnm⁵Um_34_ as C⁵-U_34_; and MnmA-dependent thiolation of C^2^-U_34_ as C^2^-U_34_. Note that for C^5^-U_34_ modification, MnmEG- and MnmC-dependent modifications could not be distinguished bioinformatically, but these are analyzed separately in the subsequent genetic screen (Fig. 2b,c).

For modifications at positions 32 and 37, codons decoded by tRNAs carrying the corresponding modification were assigned to that modification pathway. For position-34 wobble modifications, assignments were restricted to codons decoded through wobble or expanded wobble pairing by the modified anticodon, rather than to all codons readable by the same tRNA. This criterion was intended to capture codons whose decoding is most directly influenced by the wobble modification, consistent with previous studies showing that such modifications can rebalance decoding kinetics in favour of wobble over Watson–Crick pairing^1–3^. Because the relative contribution of alternative compatible isoacceptors remains unknown in vivo, two assignment models were used. In the permissive model, a codon was assigned to a modification pathway when at least one compatible tRNA carried the corresponding modification, whereas in the conservative model, assignment required all compatible tRNAs to carry it. The permissive model was used for the main analyses (Fig. 1b and Fig. 4a; Extended Data Fig. 1a,c,e,g,i), whereas the conservative model was used as a robustness analysis (Extended Data Fig. 1b,d,f,h,j,k). For each gene and modification pathway, usage of the assigned synonymous codons was normalized genome-wide as a z-score (Supplementary Data 3).

For each coding sequence and modification pathway, we calculated the relative usage of codons associated with that modification within the relevant amino-acid families. Codons assigned to a modification pathway were summed across the relevant amino-acid families and divided by the total number of codons belonging to those amino-acid families. This generated a gene-level score reflecting tRNA modification-associated synonymous codon bias within the relevant amino-acid families while excluding non-informative amino acids. For each genome and modification pathway, raw percentages were normalized across all coding sequences as z-scores by subtracting the genome-wide mean and dividing by the genome-wide standard deviation. For visualization purposes, z-scores were transformed as sign(z) × log₂(|z| + 1). Positive values therefore indicate enrichment in codons associated with a given modification pathway, in comparison to genomic average values, whereas negative values indicate depletion.

Gene-level modification-enrichment scores were compared between functional gene groups using two-sided Mann–Whitney U-tests. P values were adjusted for multiple testing using the Benjamini–Hochberg procedure across all modification pathways within each analysis. For cross-species analyses, the same scoring procedure was applied independently to *S*Tm SL1344 (GCF_000210855.2), *Shigella flexneri* 2a str. 301 (GCF_000006925.2), *E. coli* EHEC O157:H7 str. Sakai (GCF_000008865.2), *E. coli* UPEC UTI89 (GCF_000013265.1) and *Klebsiella pneumoniae* subsp. *pneumoniae* HS11286 (GCF_000240185.1). Scores were normalized independently within each genome. Virulence and ribosomal-protein genes were defined separately for each species as previously described^22^, and their score distributions were compared using two-sided Mann–Whitney U-tests followed by Benjamini–Hochberg correction. Cross-species heatmap values correspond to median(virulence genes) − median(ribosomal-protein genes), such that positive values indicate greater modification score enrichment in virulence genes.

### Mapping gene clusters in a 2D codon usage space

Genome-wide codon-usage structure was analyzed with the CodonPipe framework (DOI: 10.5281/zenodo.21303183) as described previously^22^. Briefly, relative codon usage tables were calculated for all genes of *S*Tm SL1344. Codon-usage features were centered and scaled across the entire genome (z-scores) before dimensionality reduction using UMAP, followed by k-means clustering. Functional gene clusters including ribosomal protein genes, plasmid genes, virulence genes, SPI-2 genes and genes encoding proteins depleted in the *miaA* and *mnmE* inactive-mutant proteomes were mapped into the 2D UMAP space. The latter corresponded to the 30 most depleted proteins detected in the *miaA* and *mnmE* inactive-mutant proteomes. Spatial distributions of each gene cluster were compared with the distribution of the *miaA* and *mnmE* depleted protein cluster using the Fasano–Franceschini two-dimensional Kolmogorov–Smirnov test with permutation-derived P values, followed by Benjamini–Hochberg correction for multiple comparisons.

### Conservation analysis of tRNA modification enzymes across bacterial phyla

Conservation of MiaA, MnmA, MnmE, MnmG, Tgt and TilS across bacteria was assessed using KEGG Orthology annotations. Bacterial genome representatives were retrieved from KEGG GENOME using the KEGG REST list/genome/bacteria endpoint. For each genome, the corresponding NCBI Taxonomy identifier was obtained through KEGG genome-to-taxonomy links, and bacterial phyla were assigned from NCBI Taxonomy lineages retrieved through Entrez E-utilities. Ortholog presence was defined by the presence of at least one KEGG gene assigned to the relevant KEGG Orthology group: MiaA/K00791, MnmE/K03650, MnmG/K03495, MnmA/K00566, Tgt/K00773 and TilS/K04075. For each bacterial phylum, conservation was calculated as the fraction of KEGG bacterial genome representatives assigned to that phylum that encoded each ortholog. Phyla represented by fewer than 20 KEGG bacterial genome representatives were discarded. This analysis was implemented in Python and the script is available at Zenodo (DOI: 10.5281/zenodo.21303131).

To validate the apparent depletion of canonical MnmE/MnmG annotations in Actinomycetota, representative Actinomycetota species were further queried in UniProtKB using taxonomy-restricted searches for the corresponding gene names, synonyms and protein annotations, including mnmE/trmE/thdF for MnmE and mnmG/gidA/mto1 for MnmG. UniProt hits were scored according to gene-name matches, protein-name matches and functional or domain annotations, and the best-supported hits were compared with KEGG Orthology assignments for the same species. This targeted annotation check was used to distinguish likely absence of canonical MnmE/MnmG ortholog annotations from simple KEGG assignment failure. Representative Actinomycetota species included *Mycobacterium tuberculosis*, *Mycobacterium smegmatis*, *Corynebacterium glutamicum*, *Streptomyces coelicolor*, *Streptomyces lividans*, *Nocardia farcinica* and *Bifidobacterium longum*. This analysis was implemented in Python and the script is available at Zenodo (DOI: 10.5281/zenodo.21303137).

### Cross-species comparison of synonymous codon usage

Codon counts were calculated for all coding sequences of *S*Tm SL1344 (4,868 CDS; GCF_000210855.2) and *S. aureus* NCTC 8325 (2,767 CDS; GCF_000013425.1). For the pooled codon-usage comparison in Fig. 4b and Extended Data Fig. 3b, codon counts were summed within each functional gene set and converted to relative usage within the corresponding synonymous amino-acid family. Values for ribosomal-protein, virulence and prophage genes are shown as differences from the genome-wide relative usage. For the gene-level distributions in Fig. 4c and Extended Data Fig. 3c,d, relative synonymous codon usage was calculated separately for each coding sequence; genes lacking the relevant amino acid were excluded from that codon’s analysis. Gene cluster definitions are provided in Supplementary Data 4.

## Supporting information

Supplementary Data 1. Machine-readable decoding table + literature evidence

Supplementary Data 2. Virulence cluster codon usage tables

Supplementary Data 3. tRNA modification scores across Enterobacterales

Supplementary Data 4. Gene clusters composition

Supplementary Data 5. Proteomes

Supplementary Data 6. Codon-level statistics for analysis of mutants proteomes

Supplementary Data 7. Strains and plasmids

## Data availability

All mass spectrometry proteomics data associated with this manuscript have been deposited to the ProteomeXchange consortium via MassIVE (https://massive.ucsd.edu) with the accession numbers MassIVE MSV000103162 / PRIDE PXD083883.

Genome sequences and annotations were obtained from NCBI RefSeq under accessions GCF_000210855.2 (*Salmonella enterica* serovar Typhimurium SL1344), GCF_000006925.2 (*Shigella flexneri* 2a str. 301), GCF_000008865.2 (*Escherichia coli* O157:H7 Sakai), GCF_000013265.1 (*E. coli* UTI89), GCF_000240185.1 (*Klebsiella pneumoniae* HS11286) and GCF_000013425.1 (*Staphylococcus aureus* NCTC 8325).

Source data underlying all quantitative main and Extended Data figures are provided in the Supplementary Data files indicated in their respective figure legends:

- Supplementary Data 1. Machine-readable decoding table + literature evidence
- Supplementary Data 2. Virulence cluster codon usage tables
- Supplementary Data 3. tRNA modification scores across Enterobacterales
- Supplementary Data 4. Gene clusters composition
- Supplementary Data 5. Proteomes
- Supplementary Data 6 - Codon-level statistics for analysis of mutants’ proteomes
- Supplementary Data 7. Strains and plasmids

## Code availability

Codon-usage and tRNA-enrichment analyses based on codon usage were performed using the previously described CodonPipe framework^22^, updated to version 1.1 for this study. CodonPipe v1.1 is available on Zenodo at DOI: 10.5281/zenodo.21303183

The fluorescent-reporter recoding pipeline, version 1.0, is available on Zenodo at DOI: 10.5281/zenodo.17072594

The custom Python pipeline used to analyse the conservation of tRNA modification enzymes based on KEGG Orthology annotations is available on Zenodo at DOI: 10.5281/zenodo.21303131

The custom Python pipeline used to analyse the conservation of tRNA modification enzymes based on UniProtKB annotations is available on Zenodo at DOI: 10.5281/zenodo.21303137

## Acknowledgements

We thank Anna Chirkova, Aaron Voitl, and Beatrice Claudi for help with cloning, chemostat cultures, and animal experiments.

## Author Contributions Statement

Conceptualization: FG, DB; Methodology: FG; Investigation: FG, DB, MAV; Visualization: FG; Funding acquisition: FG, DB; Project administration: FG, DB; Supervision: FG, DB; Writing – original draft: FG; Writing – review & editing: FG, DB.

## Additional information

Supplementary information is available for this paper. Correspondence and requests for materials should be addressed to Frédéric Goormaghtigh or Dirk Bumann.

## Extended data

**Extended Data Fig. 1.**
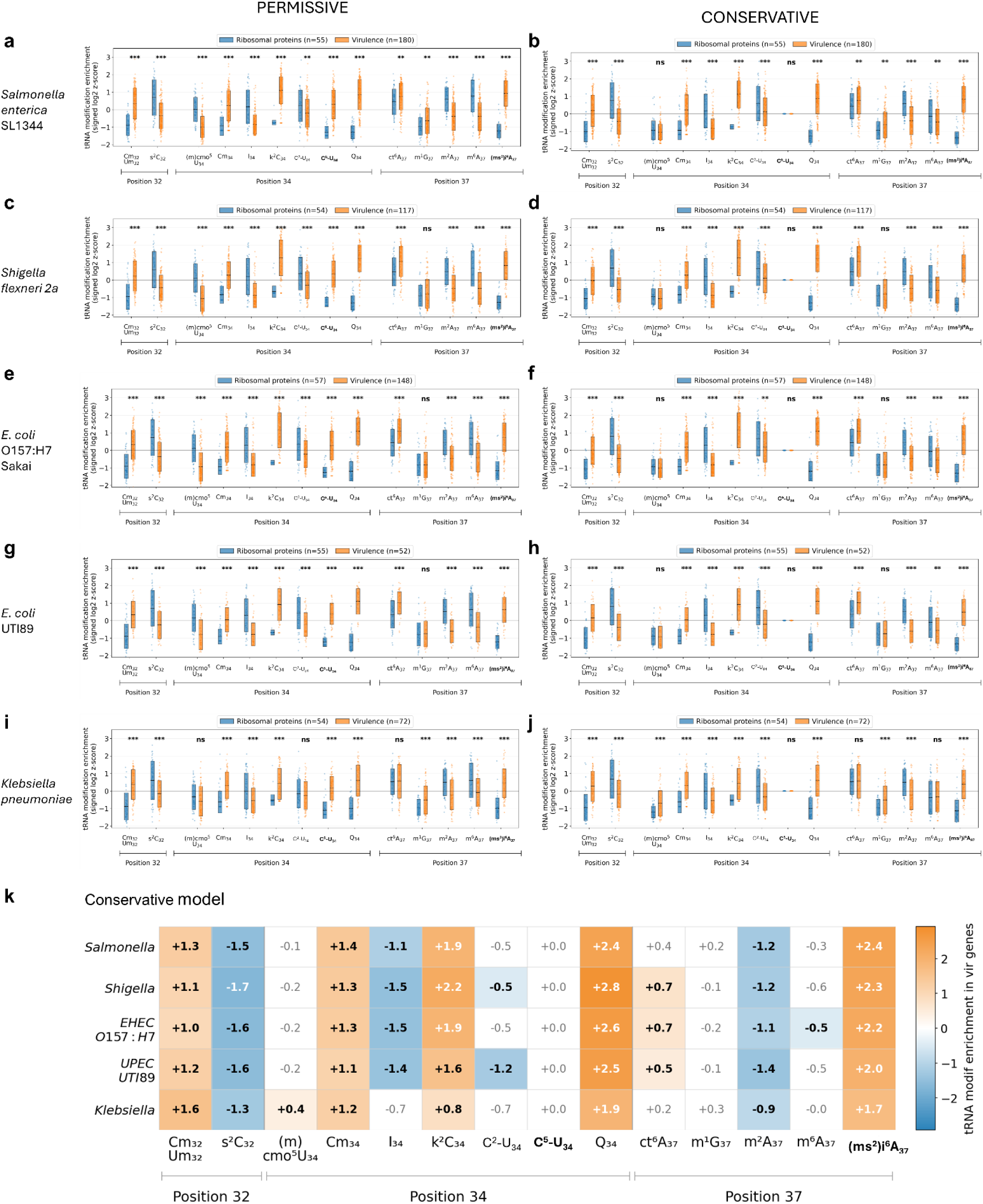
Virulence-associated tRNA modification signatures are conserved across five Enterobacterales pathogens. **a–j,** Gene-level tRNA modification enrichment scores for ribosomal-protein (blue) and virulence (orange) genes using permissive (left) and conservative (right) codon-assignment models. The permissive model assigns a codon when at least one compatible decoder carries the modification, whereas the conservative model requires all compatible decoders to carry it. Scores are shown for *Salmonella enterica* SL1344 (**a**,**b**), *Shigella flexneri 2a* (**c**,**d**), *E. coli* EHEC O157 Sakai (**e**,**f**), *E. coli* UPEC UTI89 (**g**,**h**) and *Klebsiella pneumoniae* (**i**,**j**). Each dot represents one gene. Scores were normalized genome-wide as z-scores and transformed as sign(z) × log₂(|z| + 1). Statistical significance was assessed using two-sided Mann–Whitney U-tests with Benjamini–Hochberg correction across modification comparisons within each analysis. **k**, Cross-species comparison using the conservative model. Values indicate the median score difference between virulence and ribosomal-protein genes; orange indicates higher scores in virulence genes and blue higher scores in ribosomal-protein genes. Bold values indicate q < 0.001 after Benjamini–Hochberg correction. White cells are not significant. Gene cluster definitions are provided in Supplementary Data 4 and tRNA modification scores are provided in Supplementary Data 3.

**Extended Data Fig. 2.**
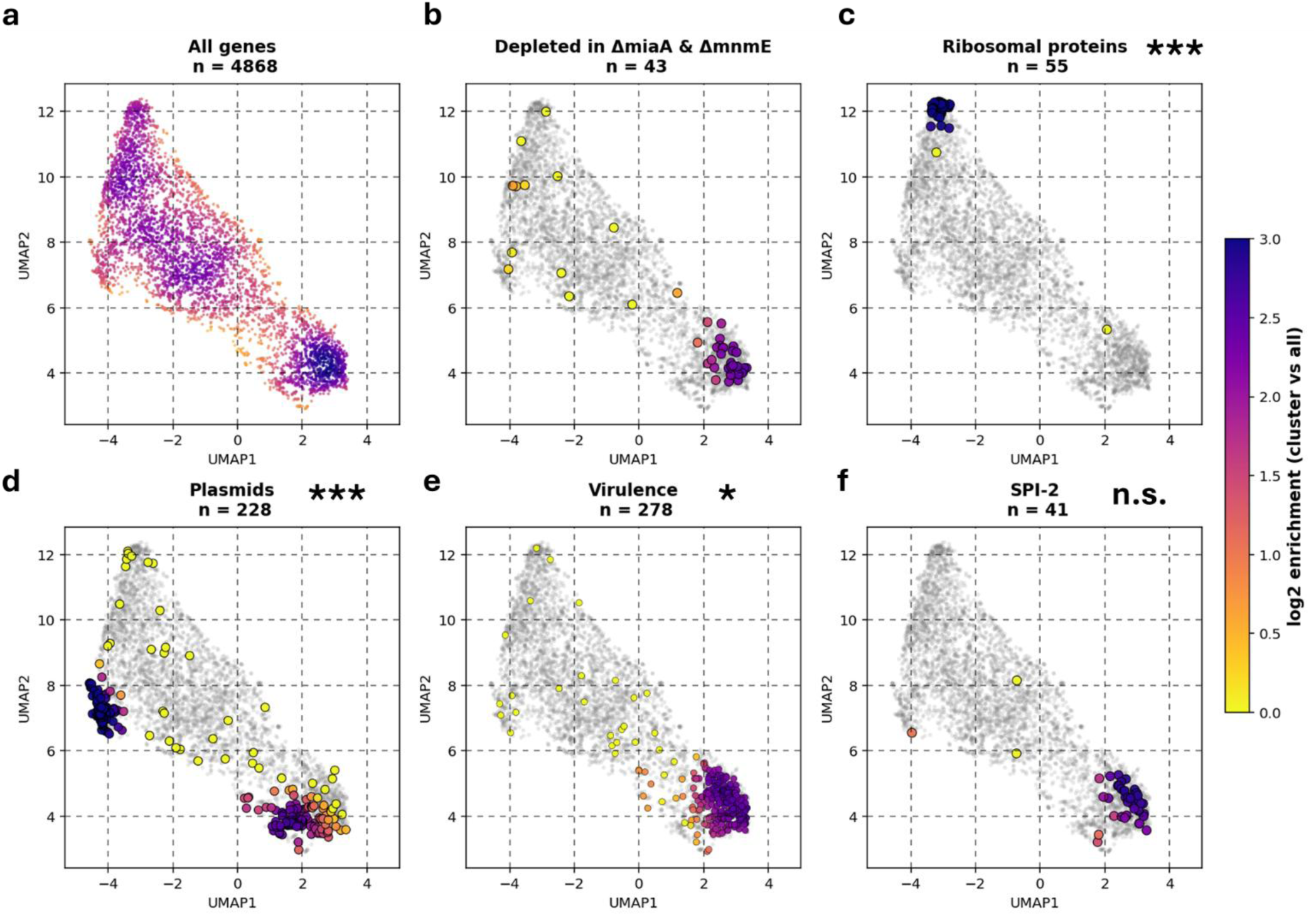
Proteins depleted in *miaA\** and *mnmE\** occupy a virulence-associated codon-usage space. **a**, UMAP projection of all *STm* coding sequences based on codon-usage profiles, followed by k-means clustering as described previously^22^. **b–f**, Spatial enrichment of the union of the 30 most depleted proteins in each of the *miaA\** and *mnmE\** proteomes (**b**), ribosomal-protein genes (**c**), plasmid genes (**d**), virulence genes (**e**) and SPI-2 genes (**f**) within the same codon-usage landscape. Grey points indicate all coding sequences and colored overlays indicate local enrichment of each gene set relative to the whole-genome background. Color intensity represents log_2_ enrichment. Gene counts are indicated above each panel. Spatial distributions were compared with the depleted-protein set using the Fasano–Franceschini two-dimensional Kolmogorov–Smirnov test with Benjamini–Hochberg correction (*q < 0.05, **q < 0.01, ***q < 0.001, n.s., not significant). Gene cluster definitions are provided in Supplementary Data 4.

**Extended Data Fig. 3.**
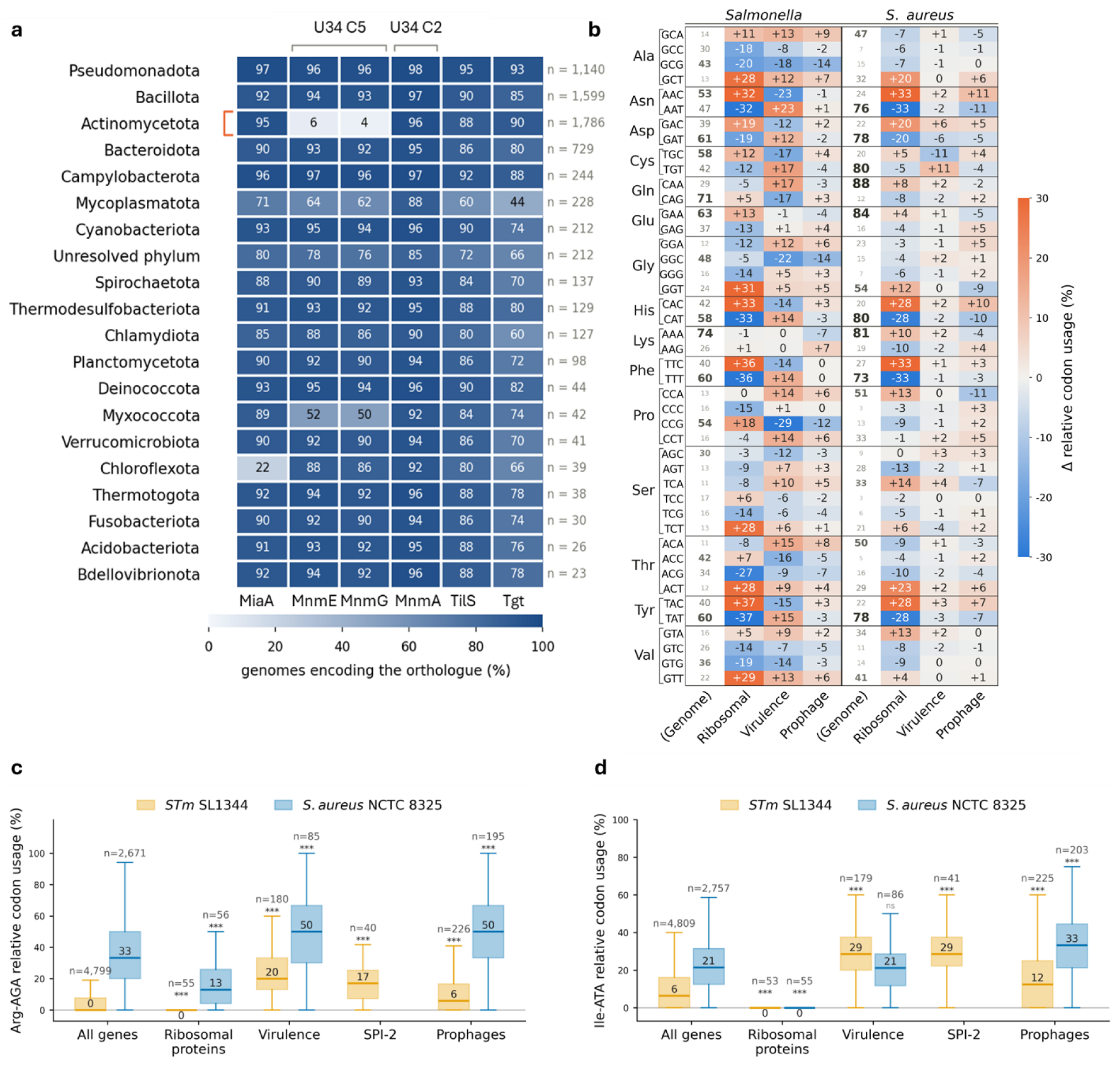
tRNA modifications are broadly conserved, but their role could be rewired in distant AT-rich pathogens, when codon usage changes dramatically. **a**, Conservation of major tRNA modification enzymes across bacterial phyla. Cells indicate the percentage of KEGG bacterial genomes containing at least one orthologue of MiaA, MnmE, MnmG, MnmA, TilS or Tgt. n indicates the number of genomes analyzed per phylum. Only phyla represented by at least 20 genomes are shown. The red bracket highlights the low representation of canonical MnmE/MnmG orthologues in Actinomycetota. **b**, Synonymous codon usage in *S*Tm SL1344 and *S. aureus* NCTC 8325. Genome values indicate relative usage within each synonymous codon family (%), with font size scaled to frequency. Ribosomal, virulence and prophage columns show differences from the genome-wide average (blue, depleted, orange, enriched). The 44 synonymous codons that are not shown in Fig. 4b are displayed here; the 15 leucine, arginine and isoleucine codons are in Fig. 4b. **c**,**d**, Gene-level relative usage of Arg-AGA (**c**) and Ile-ATA (**d**) across functional gene classes in *S*Tm and *S. aureus*. Numbers above groups indicate gene counts and values within boxes indicate median codon usage (%). SPI-2 is specific to *S*Tm and therefore has no corresponding *S. aureus* group. Statistical significance was assessed using two-sided Mann–Whitney U-tests; n.s., not significant, ***q < 0.001. Gene cluster definitions are provided in Supplementary Data 4.

**Extended Data Table 1.**
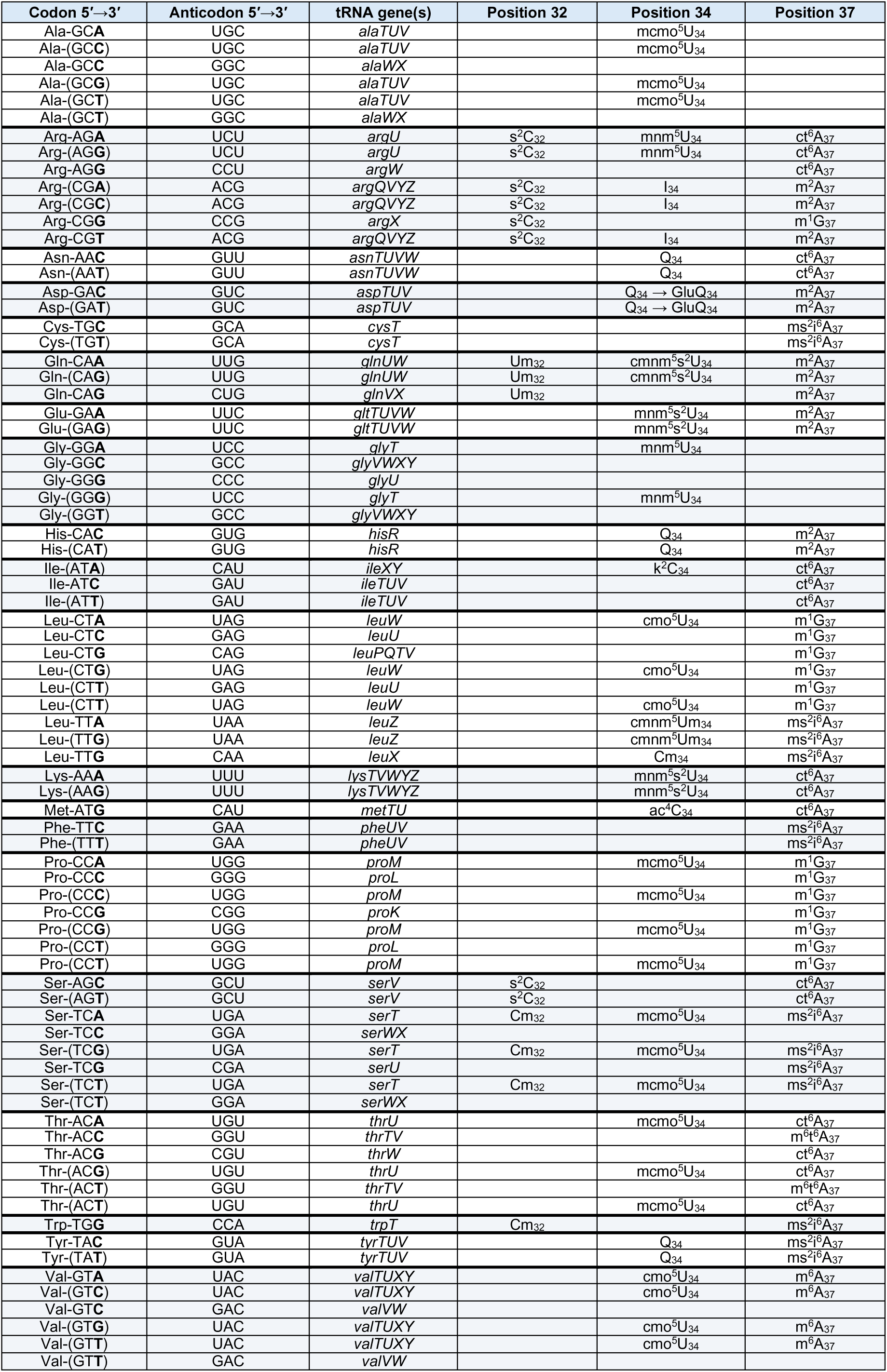
Curated codon-anticodon-tRNA modification decoding table. Codons and anticodons are shown 5′ to 3′. Each row lists a codon, a compatible decoder anticodon, the corresponding tRNA species and curated modifications at positions 32, 34 and 37. Codons in parentheses indicate wobble or expanded-decoding assignments. Assignments were curated from MODOMICS^25^, available reviews^7^ and pathway-specific literature, including Cm_32_/Um ^29^, s^2^C_32_ ^72^, C^2^- and C^5^-U_34_ modifications^1,46^, Cm_34_/cmnm^5^Um ^28^, GluQ_34_/Q_34_ ^73^, (m)cmo^5^U_34_ ^2^, m^2^A ^74^, (ms^2^)i^6^A ^13,40^, I_34_ ^75^, k^2^C_34_ ^76^, ac^4^C_34_ ^77^, m^1^G_37_ ^78^, ct^6^A ^79,80^, m^6^t^6^A ^81^ and m^6^A ^82^. Related modifications were grouped into pathway-level classes for tRNA modification enrichment analyses as described in Methods. A more detailed and machine-readable table together with exhaustive literature evidence and description is provided in Supplementary Data 1.

**Extended Data Table 2.** Grouping of individual tRNA modifications into pathway classes. The modifications listed in Extended Data Table 1 were grouped into fifteen classes, each corresponding to one biosynthetic route. These classes are the functional units used in Fig. 1b, Fig. 4a, Extended Data Fig. 1 and Supplementary Data 3. cmnm^5^Um_34_ appears in two classes because it carries both a TrmL-dependent 2′-O-methyl group and an MnmEG-dependent C^5^ side chain. C^5^-U_34_ and C^2^-U_34_ distinguish dependence on MnmEG alone from dependence on MnmEG together with MnmA. ac^4^C_34_ was curated but not scored (Methods).

| Pathway class | Individual modifications grouped into it | Enzymes |
| --- | --- | --- |
| Cm<sub>32</sub>/Um<sub>32</sub> | Cm<sub>32</sub>, Um<sub>32</sub> | TrmJ |
| s<sup>2</sup>C<sub>32</sub> | s<sup>2</sup>C<sub>32</sub> | TtcA |
| (m)cmo<sup>5</sup>U<sub>34</sub> | cmo<sup>5</sup>U<sub>34</sub>, mcmo<sup>5</sup>U<sub>34</sub> | CmoA, CmoB, CmoM |
| Cm<sub>34</sub> | Cm<sub>34</sub>, cmnm<sup>5</sup>Um<sub>34</sub> | MnmE, MnmG, TrmL |
| I<sub>34</sub> | I<sub>34</sub> | TadA |
| k<sup>2</sup>C<sub>34</sub> | k<sup>2</sup>C<sub>34</sub> | TilS |
| ac<sup>4</sup>C<sub>34</sub> | ac<sup>4</sup>C<sub>34</sub> (curated, not scored) | TmcA |
| Q<sub>34</sub> | Q<sub>34</sub>, GluQ<sub>34</sub> | Tgt, QueA, QueG, GluQ-RS |
| C<sup>5</sup>-U<sub>34</sub> | cmnm<sup>5</sup>U<sub>34</sub>, nm<sup>5</sup>U<sub>34</sub>, mnm<sup>5</sup>U<sub>34</sub>, cmnm<sup>5</sup>Um<sub>34</sub> | MnmE, MnmG, MnmC |
| C<sup>2</sup>-U<sub>34</sub> | cmnm<sup>5</sup>s<sup>2</sup>U<sub>34</sub>, mnm<sup>5</sup>s<sup>2</sup>U<sub>34</sub> | MnmA |
| ct<sup>6</sup>A<sub>37</sub> | ct<sup>6</sup>A<sub>37</sub>, m<sup>6</sup>t<sup>6</sup>A<sub>37</sub> | TsaB-E, TcdA, TrmO |
| m<sup>1</sup>G<sub>37</sub> | m<sup>1</sup>G<sub>37</sub> | TrmD |
| m<sup>2</sup>A<sub>37</sub> | m<sup>2</sup>A<sub>37</sub> | RlmN |
| m<sup>6</sup>A<sub>37</sub> | m<sup>6</sup>A<sub>37</sub> | YfiC (TrmM) |
| (ms<sup>2</sup>)i<sup>6</sup>A<sub>37</sub> | i<sup>6</sup>A<sub>37</sub>, ms<sup>2</sup>i<sup>6</sup>A<sub>37</sub>, ms<sup>2</sup>io<sup>6</sup>A<sub>37</sub> | MiaA, MiaB, MiaE |

**Extended Data Table 3.**
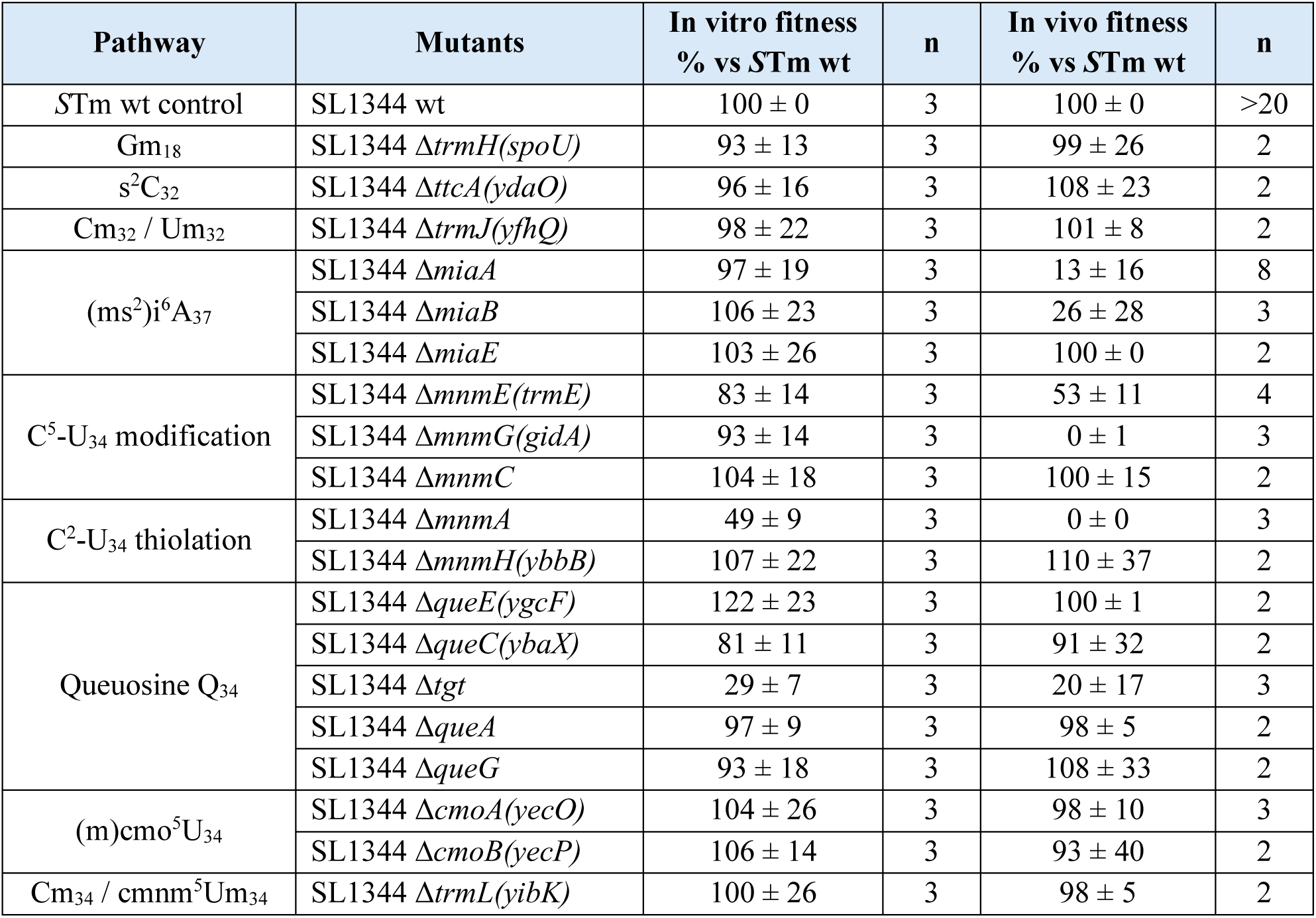
*In vitro* and *in vivo* fitness of tRNA modification enzyme deletion mutants. Deletion mutants affecting anticodon-loop tRNA modification pathways were screened for fitness defects *in vitro* and during systemic infection. *In vitro* fitness was calculated from growth rates in MES-ch medium and expressed relative to WT. For *in vivo* fitness, BALB/c mice were co-infected with WT and fluorescently distinguishable isogenic mutants. Bacterial divisions after four days of infection were estimated from the change in splenic bacterial abundance and mutant fitness was expressed relative to the co-infected WT. Values are mean ± s.d. n indicates independent biological replicates for *in vitro* measurements and individual mice for *in vivo* measurements.

**Extended Data Table 4.** Catalytically inactive tRNA modification mutants used in this study. Alleles were selected based on residues or motifs previously shown to be required for catalytic activity, substrate recognition or cofactor binding of the corresponding tRNA modification enzymes. All listed alleles were obtained and analyzed except *mnmG\**, which could not be recovered despite repeated attempts, as described in the Methods.

| Mutants | Inactive alleles | Function disrupted | Refs |
| --- | --- | --- | --- |
| *miaA* * | T19A K23A | Residues required for DMAPP-dependent catalysis and substrate binding | 56,57 |
| *miaB* * | C157A C161A C164A | Iron–sulfur cluster motif | 58 |
| *mnmA* * | Q151E | Recognition of the U<sub>34</sub> nucleotide in target tRNAs | 59 |
| *mnmE* * | G232A | GTPase activity | 60 |
| *mnmG* * | G13A G15A | Rossmann-fold FAD-binding motif | 61,83 |
| *tgt* * | C302A C304A C307A | Zinc-binding subdomain required for tRNA anchoring | 62 |

**Extended Data Table 5.** Functional enrichment analysis of catalytically inactive tRNA modification mutants. For each mutant and functional category, the table reports the number of quantified proteins (n), median log_2_FC, and the number of proteins with negative FC. Directional bias was assessed using a one-sided exact binomial sign test, and category-level abundance shifts using a two-sided Mann–Whitney U-test against all other quantified proteins in the same comparison. P values were corrected for multiple testing using Benjamini–Hochberg procedure across the 13 categories within each mutant, separately for each test. Q values below 0.05 are shown in bold. Category assignments are provided in Supplementary Data 4. Proteins may belong to more than one category, and categories with fewer than five quantified proteins were not tested. Categories appear in the same order in all four panels. FC, fold-change.

| Cluster | *mnme\** (G232A) |  |  |  |  | *miaA\** (T19A/K23A) |  |  |  |  |
| --- | --- | --- | --- | --- | --- | --- | --- | --- | --- | --- |
|  | n | Median log<sub>2</sub>FC | Neg. | Sign *q* | MWU *q* | n | Median log<sub>2</sub>FC | Neg. | Sign *q* | MWU *q* |
| SPI-2 + effectors | 24 | -2.57 | 24/24 | **4 × 10<sup>-7</sup>** | **3 × 10<sup>-15</sup>** | 24 | -1.48 | 23/24 | **1 × 10<sup>-5</sup>** | **1 × 10<sup>-11</sup>** |
| Virulence (all) | 62 | -1.21 | 53/62 | **7 × 10<sup>-8</sup>** | **7 × 10<sup>-19</sup>** | 61 | -1.04 | 55/61 | **3 × 10<sup>-10</sup>** | **5 × 10<sup>-20</sup>** |
| SPI-1 | 6 | 0.35 | 2/6 | 1 | 0.92 | 5 | -0.39 | 4/5 | 0.24 | 0.18 |
| O-antigen | 23 | -0.23 | 14/23 | 0.53 | 0.11 | 23 | -0.29 | 19/23 | **0.0042** | **0.026** |
| Prophages | 15 | -0.65 | 13/15 | **0.012** | **0.0008** | 14 | -0.52 | 11/14 | **0.046** | **0.034** |
| Plasmids | 24 | -0.22 | 14/24 | 0.59 | 0.11 | 24 | -0.24 | 17/24 | **0.046** | 0.16 |
| Ribosomal proteins | 49 | -0.29 | 36/49 | **0.003** | **0.0044** | 49 | 0.13 | 4/49 | 1 | **2 × 10<sup>-7</sup>** |
| tRNA charging | 21 | 0.29 | 7/21 | 1 | 0.84 | 21 | 0.33 | 3/21 | 1 | **4 × 10<sup>-5</sup>** |
| ABC transporters | 82 | 0.28 | 27/82 | 1 | **0.014** | 80 | -0.34 | 60/80 | **2 × 10<sup>-5</sup>** | **1 × 10<sup>-5</sup>** |
| C metabolism | 97 | 0.1 | 45/97 | 1 | 0.85 | 96 | -0.19 | 60/96 | **0.02** | 0.23 |
| Respiration | 43 | 0.01 | 21/43 | 0.95 | 0.84 | 41 | -0.23 | 27/41 | **0.046** | 0.33 |
| Cytochromes | 6 | 0.13 | 3/6 | 0.95 | 0.98 | 6 | 0.35 | 2/6 | 1 | 0.16 |
| Two-component system | 29 | -0.05 | 15/29 | 0.93 | 0.15 | 29 | -0.20 | 22/29 | **0.011** | 0.072 |

**Extended Data Table 5 | Functional enrichment analysis of catalytically inactive tRNA modification mutants.**
| Cluster | *tgt\** (C302A/C304A/C307A) |  |  |  |  | *miaB\** (C157A/C161A/C164A) |  |  |  |  |
| --- | --- | --- | --- | --- | --- | --- | --- | --- | --- | --- |
|  | n | Median log<sub>2</sub>FC | Neg. | Sign *q* | MWU *q* | n | Median log<sub>2</sub>FC | Neg. | Sign *q* | MWU *q* |
| SPI-2 + effectors | 30 | -0.58 | 27/30 | **3 × 10<sup>-5</sup>** | **1 × 10<sup>-10</sup>** | 31 | -0.08 | 19/31 | 0.61 | **0.012** |
| Virulence (all) | 69 | -0.46 | 62/69 | **3 × 10<sup>-11</sup>** | **2 × 10<sup>-16</sup>** | 72 | -0.05 | 40/72 | 0.67 | **0.012** |
| SPI-1 | 6 | -0.01 | 3/6 | 0.71 | 0.95 | 6 | 0.03 | 3/6 | 1 | 0.87 |
| O-antigen | 23 | -0.09 | 15/23 | 0.17 | 0.53 | 23 | -0.02 | 13/23 | 0.88 | 0.21 |
| Prophages | 13 | -0.20 | 10/13 | 0.11 | **0.039** | 14 | -0.07 | 10/14 | 0.58 | **0.021** |
| Plasmids | 24 | -0.07 | 13/24 | 0.5 | 0.95 | 25 | 0.02 | 11/25 | 1 | 0.54 |
| Ribosomal proteins | 49 | 0.02 | 23/49 | 0.72 | 0.28 | 49 | -0.07 | 35/49 | **0.025** | **0.0001** |
| tRNA charging | 21 | -0.03 | 15/21 | 0.11 | 0.95 | 21 | 0.03 | 10/21 | 1 | 0.54 |
| ABC transporters | 84 | -0.01 | 45/84 | 0.42 | 0.28 | 84 | 0.13 | 32/84 | 1 | 0.35 |
| C metabolism | 97 | -0.03 | 57/97 | 0.11 | 0.39 | 97 | 0.08 | 34/97 | 1 | 0.54 |
| Respiration | 41 | -0.14 | 32/41 | **0.0009** | **0.0084** | 42 | 0.12 | 11/42 | 1 | 0.54 |
| Cytochromes | 6 | -0.02 | 4/6 | 0.45 | 0.95 | 6 | 0.15 | 1/6 | 1 | 0.55 |
| Two-component system | 30 | -0.09 | 19/30 | 0.17 | 0.28 | 30 | 0.15 | 8/30 | 1 | 0.39 |

**Extended Data Table 6.** Previous studies linking tRNA modification and codon-dependent translation to bacterial virulence. Published studies reporting effects of tRNA-modification perturbation or tRNA reprogramming on bacterial virulence, virulence-gene expression or infection fitness are summarized. For each study, the table lists the affected gene or pathway, bacterial species, experimental infection model, principal phenotype and reference.

| Genes | Bacterial species | Infection model | Description | Ref |
| --- | --- | --- | --- | --- |
| *miaA* | *E. coli* UPEC UTI89 | BALB/c mice | UTI89 Δ*miaA* was outcompeted by WT UTI89 during gastrointestinal colonization of BALB/c mice. | 13 |
| *mnmE*<br>*mnmG* | *Shigella flexneri* | Gut epithelial organoid model | MnmE/G-dependent tRNA modification promotes expression of *pINV* virulence genes enriched in modification-sensitive codons, including AGA-Arg, AGG-Arg, GGA-Gly. | 10 |
| *mnmG* | *Pseudomonas aeruginosa* | *Galleria mellonella*<br>BALB/c mice | Loss of *GidA*/*MnmG* impaired virulence in *G. mellonella* and competitive fitness in vivo and altered translation and expression of virulence-associated genes. | 14 |
| *miaA* | *Salmonella* Typhimurium | Immunodeficient gp91-/- phox and C57BL/6 WT mice | *miaA* disruption caused attenuation during systemic infection. | 11 |
| *mnmE*,<br>*mnmG* | *Salmonella* Typhimurium | Swiss-Webster mice | Δ*mnmE* and Δ*mnmG* mutants showed impaired invasion and intracellular replication and were strongly attenuated in mice. | 12 |
| *miaA* | *Salmonella* Typhimurium | BALB/c mice | *miaA* was identified as required for full fitness during systemic infection. | 8 |
| *mnmE*,<br>*mnmG* | *Streptococcus pyogenes* | SKH1 hairless mice, subcutaneous soft-tissue infection | MnmG and MnmE control expression of multiple virulence factors; *GidA*-deficient strains were strongly attenuated in a murine soft-tissue infection model. | 42 |
| *miaA* | *P. aeruginosa* | *Galleria mellonella* | Loss of *MiaA* impaired decoding of *MiaA*-sensitive codons, broadly altered virulence-associated gene expression and reduced pathogenicity in *G. mellonella*. | 15 |
| - | *E. coli* EHEC O157:H7 | In vitro, norepinephrine-induced virulence conditions | Norepinephrine remodeled tRNA modification and abundance and shifted protein synthesis toward A/U-ending codon-biased virulence genes, including the LEE pathogenicity island. | 16 |
| *miaA*, *tgt* | *Shigella flexneri* | HeLa-cell invasion/focus-plaque and in-vitro virulence assays | Loss of *MiaA* or *Tgt* reduced *VirF* abundance and virulence. *VirF* overexpression rescued the attenuated *tgt* and avirulent *miaA* phenotypes, identifying *VirF* translation as a major modification-sensitive regulatory node. | 18 |
| *mnmA* | *Mycobacterium tuberculosis* | Macrophage intracellular-growth model | Loss of *MnmA* abolished *MnmA*-dependent U34 thiolation and impaired intracellular growth in macrophages. | 84 |
| *mnmE* | *Francisella tularensis* | Macrophage/intracellular growth | *mnmE* disruption reduced *Francisella* pathogenicity-island expression and intracellular growth and altered the ppGpp-dependent regulatory network controlling virulence genes. | 85 |
| *tgt* | *Vibrio cholerae* | Sub-MIC aminoglycoside stress | *Tgt*-dependent Q<sub>34</sub> modification altered decoding of Tyr-TAT/TAC codons and selectively modulated translation of codon-biased transcripts. | 86 |

